# Type two secretion systems secretins are necessary for exopolymeric slime secretion in cyanobacteria and myxobacteria

**DOI:** 10.1101/2022.03.08.483542

**Authors:** David M. Zuckerman, Jeffery Man To So, Egbert Hoiczyk

## Abstract

While protein translocation in Gram-negative bacteria is well understood, our knowledge about the translocation of other high-molecular-weight substances is limited. Nozzle-like structures that secrete exopolymeric substances during gliding motility have previously been observed in the outer membranes of cyanobacteria and myxobacteria. Here, we show that these nozzles are composed of the secretins PilQ/GspD, the outer membrane component of the type II and III secretion systems, the type IV pilus apparatus, and filamentous phage extrusion machinery. Our results show for the first time that secretins may be used for secretion of non-proteinaceous polymers in some bacteria, considerably expanding the repertoire of substrates of these multifunctional outer membrane gates. Moreover, we show that *gspD* is an essential gene in *Myxococcus xanthus*, which, when depleted, renders this bacterium defective in slime secretion and gliding motility.

**Significance:** Many bacteria exhibit gliding motility, movement across surfaces. This motility has been correlated with the deposit of slime trails in their wake. To date, the mechanism of slime secretion has not been understood, and no cell envelope-structures have been identified that are involved in slime secretion during gliding motility. Here, we show that cyanobacteria and myxobacteria use the secretins PilQ/GspD, the outer membrane channels of the T2SS, for slime secretion, which demonstrates a novel cargo transport capacity of these multifunctional outer membrane gates.

## Introduction

Secretion of macromolecules is an important component of environmental adaption, and a key property of any living cell. Like eukaryotic cells, bacteria contain dedicated macromolecular secretion systems in the cell envelope that are used to translocate proteins, nucleic acids, and carbohydrates (1). Despite an extraordinary diversity of both substrates and bacterial cell physiologies, there are only a limited number of secretion systems. In Gram-negative bacteria, twelve protein (type I-IX, the Bam and Lpt machineries, and the chaperone/usher pathway (1–5)), one nucleic acid (type IV; **6**), and three carbohydrate (Wzx/Wzy, ABC transporter, and synthase-dependent (7)) secretion systems have been described. Moreover, a close inspection of these molecular machines reveals the utilization of multiple homologous proteins, suggesting divergence from common ancestry. Diversity between the systems appears to have evolved through use of novel proteins, and “mixing-and-matching” of protein components between translocation machineries (8).

One well-studied component of secretory machinery shared between several systems is the secretin family of proteins (9–11). These multimeric proteins form the outer membrane (OM) gates of the type II and III secretion systems, the type IV pilus apparatus, and filamentous phage extrusion machinery (12–14). Secretins form a functional channel with an OM pore that is 5–8 nm wide, allowing the passage of large cargo molecules such as folded proteins and multimeric protein fibers. These channels are typically formed by the assembly of 15 monomers of GspD (e.g. GspD_Ecol_ 15mer PDB ID code 5WQ7), or 12 to 14 (in some cases up to 19; **15**) monomers of PilQ (e.g. PilQ_Mxan_ 12mer PDB ID code 3JC9; PilQ_Paer_ 14mer PDB ID code 6VE3; **10, 16-18**). Nearly all secretins (with the known exception of HxcQ of *Pseudomonas aeruginosa*; **19**) require additional proteins, called pilotins or accessory proteins, for their assembly. These proteins contribute to stability, OM targeting, and oligomerization of the secretins (20). Secretins can be identified by their highly-conserved secretin domains located at or near the C-terminus of the protein, which form the OM-embedded portion of the complex (21–22). The N-terminal domains have greater variability, and create multiple ring structures that form a large periplasmic vestibule (21, 23). These N-terminal domains also form a prominent constriction between the actual secretin gate and the vestibule, termed the periplasmic gate, whose functional significance is currently not completely understood (22, 23–25). The N-terminal periplasmic domains interact with additional proteins, including the cargo and the cytoplasmic membrane-embedded platform of the secretion machinery, to facilitate the opening of the channel and the docking and release of cargo. Importantly, this process must be highly controlled to prevent unintended breaches of the OM. Although the cargos of the best-understood systems are folded proteins or protein fibers, the transient interactions with the channel should theoretically enable secretins to translocate highly diverse molecules, including non-proteinaceous ones.

While protein secretion has long been studied in Gram-negative bacteria, our understanding of the secretion of additional extracellular materials is less complete (26–28). In part, this is because the complexity of extracellular polymeric substances (EPS) is confounding. For example, many EPS species are composed of polysaccharides (29), and bacteria utilize a greater diversity of monosaccharides than amino acids. These monosaccharides are connected by various chemical linkages, and are further diversified by chemical alterations introduced by enzymatic modification (for examples see **30-31**). Despite these differences, secretion of both protein and EPS pose similar challenges to the cell, as bulky, hydrophilic, high-molecular-weight polymers are translocated across hydrophobic membranes. So far, three different mechanisms have been described for EPS secretion in Gram-negative bacteria (**28,** for a review in *Myxococcus xanthus* see **32**): the Wzx/Wzy, the ABC transporter, and the synthase-dependent secretion pathways. The Wzx/Wzy pathway is used by bacteria for the synthesis of group I capsular exopolysaccharide, O-antigen lipopolysaccharide (LPS) and succinoglycan EPS, which are synthesized from sugar phosphates that bind to a carrier lipid in the cytoplasmic membrane (26–28). Upon binding, the monomers form short oligosaccharides that are flipped across the membrane, polymerized by a periplasmic enzyme (Wzy), and fed into the Wza channel (33). The ABC transporter pathway is used for group 2 capsular polysaccharides, the LPS common antigens and N-glycosylation of outer membrane and periplasmic proteins, in which the entire carbohydrate is synthesized on a carrier lipid before being transported across the cytoplasmic membrane *via* an ABC transporter (28, 34). Both the Wzx/Wzy and the ABC transporter pathways rely on proteins of the polysaccharide co-polymerase (PCP) and OM polysaccharide export (OPX) protein families for OM translocation (35–37). Although members of the OPX protein families can be easily identified using bioinformatics, structural data for is protein families are scarce.

The only exception is the Wza channel (PDB ID code 2J58) of the Wzx/Wzy system from *E. coli* which has been resolved at atomic resolution (38). Of note, the tandem β-grasp fold that forms the periplasmic domain of Wza can also be found in the group 4 polysaccharide capsule protein GfcC (39). However, the exact role of GfcC in polymer secretion is yet to be determined (40). For the PCP protein family, full-length structures of Class 1 PCP Wzz (PDB ID code 6RBG) and Class 2 PCP Wzc (PDB ID code 7NHR) have recently been solved using cryo-electron microscopy (41, 42). The third EPS secretion mechanism, which appears to be used by bacteria for the secretion of many high-molecular-weight polysaccharide moieties, such as cellulose (43), alginate (44), and poly-*β*-D-*N*-acetylglucosamine (PNAG; **45**), is called the synthase-dependent pathway (7), referring to the fact that a cytoplasmic membrane-embedded glycosyl transferase simultaneously facilitates polymerization and trans-membrane translocation (46). Depending on the substrate in question, these steps can be performed with or without participation of a carrier lipid and, in some cases, are stimulated by the bacterial second messenger bis-(3’-5′)-cyclic dimeric guanosine monophosphate (c-di-GMP; **47**). Once in the periplasm, the newly formed polymer interacts with a tetratricopeptide repeat (TPR-) containing protein (48) and is released through an OM porin like AlgE (**49**, PDB ID code 3RBH).

While some EPS polymers with relevance to medicine and industry have been widely studied (27), the majority of EPS molecules produced by environmental bacteria are poorly characterized. One such environmental EPS, often referred to as slime, is deposited as trails behind certain gliding bacteria (50), including cyanobacteria (51) and myxobacteria (52). Although it is generally accepted that slime secretion in these organisms is important for motility (53), the precise contribution in some gliding microbes is less clear (54) due to the absence of information on the characteristics of slime. Namely, the composition of the slime, enzymes that synthesize the slime, and the slime secretion apparatus have yet to be determined.

In this study, we use structural and biochemical assays to identify the OM secretion channel for slime. We found that the secretins PilQ and GspD constitute the slime-secretion nozzles in cyanobacteria and myxobacteria, respectively. Our results show for the first time that secretins can facilitate translocation of molecules other than proteins or protein fibers, considerably expanding the repertoire of substrates of these multifunctional OM gates.

Moreover, our results show that *gspD* is an essential gene in *M. xanthus* that, when depleted, renders this bacterium defective in slime secretion and motility, confirming that GspD-facilitated secretion is essential for gliding in this bacterium. Our results add to our knowledge that secretins are involved in the secretion of toxins and pilus-mediated host attachment, finding that they also contribute to motility and potentially the formation of biofilms through exopolysaccharide secretion.

## Results

### PilQ forms the Slime Nozzle in Filamentous Cyanobacteria

Previously, we demonstrated that cyanobacteria of the genera *Oscillatoria*, *Phormidium*, *Lyngbya*, and *Anabaena* used rows of tilted nozzles (“junctional pore complexes”) at the cross walls of their multicellular filaments to secrete bands of slime (51, 55). Since these bands elongated at the same rate with which the filaments were moving, it was proposed that slime secretion powers gliding motility (56). We wished to identify the slime secretion apparatus, however, the complex culture requirements of these species made isolating these nozzles impossible at the time (57). Therefore, we initially used the more easily cultivated species *Arthrospira* (*Spirulina*) *platensis* for the current study (58). As this free-floating species is usually cultivated in aerated reactor vessels, most available clones are non- or temporarily non-motile. For that reason, we initially confirmed that our clone secreted slime using direct observations (51) and was able to glide in an established clumping assay (**59-60**; *SI Appendix*, Fig. S1). Next, thin sections of cryo-substituted cells were analyzed by electron microscopy to confirm the presence of the tilted trans-peptidoglycan channels harboring the nozzle apparatus (**Fig. 1 *A-C***). Rotary shadowing and negative staining of preparations of isolated OMs were used to directly visualize rows of nozzles (**Fig. 1*D***). Together, these results documented that the cell envelope architecture and arrangement of nozzles in *A. platensis* is identical to all of our previously studied filamentous cyanobacteria (55). To identify the major component(s) of the nozzles, we next purified cell envelopes, fractionated, and screened for the presence of nozzle-like complexes using electron microscopy. This strategy yielded nozzle-enriched fractions, devoid of any other large-scale complexes (**Fig. 1*E***). Ring-shaped top views of the complexes were also observed upon adsorption to grids without glow discharge, likely due to a preferential adsorption of the complex on these grids (**Fig. 1*F***), as previously reported (51). These nozzle-enriched fractions were separated by SDS-PAGE, and revealed two prominent protein bands at >250 and 30 kDa (**Fig. 1*G***). Mass spectrometry and Edman degradation identified these proteins as the secretin PilQ (*SI Appendix*, Table S1) and the pentapeptide repeat protein NIES39_A07680 (61). To further verify that PilQ forms the nozzles complexes, isolated nozzles were labeled using antisera raised against GspD from *M. xanthus* (see below) that cross-reacts with PilQ from *A. platensis* (*SI Appendix*, Fig. S2*A*), and visualized by immunogold labeling and electron microscopy. Anti-GspD antisera labeled about 50% of nozzles (**Fig. 1*H***), while control antisera labeled only 15% of the complexes. Finally, we averaged negatively stained *A. platensis* nozzle complexes and compared them with published averages of other secretin complexes revealing strong structural similarities even with distantly related complexes furthermore supporting our interpretation that PilQ forms the nozzles of filamentous cyanobacteria (*SI Appendix*, Fig. S3).

**Fig. 1.**
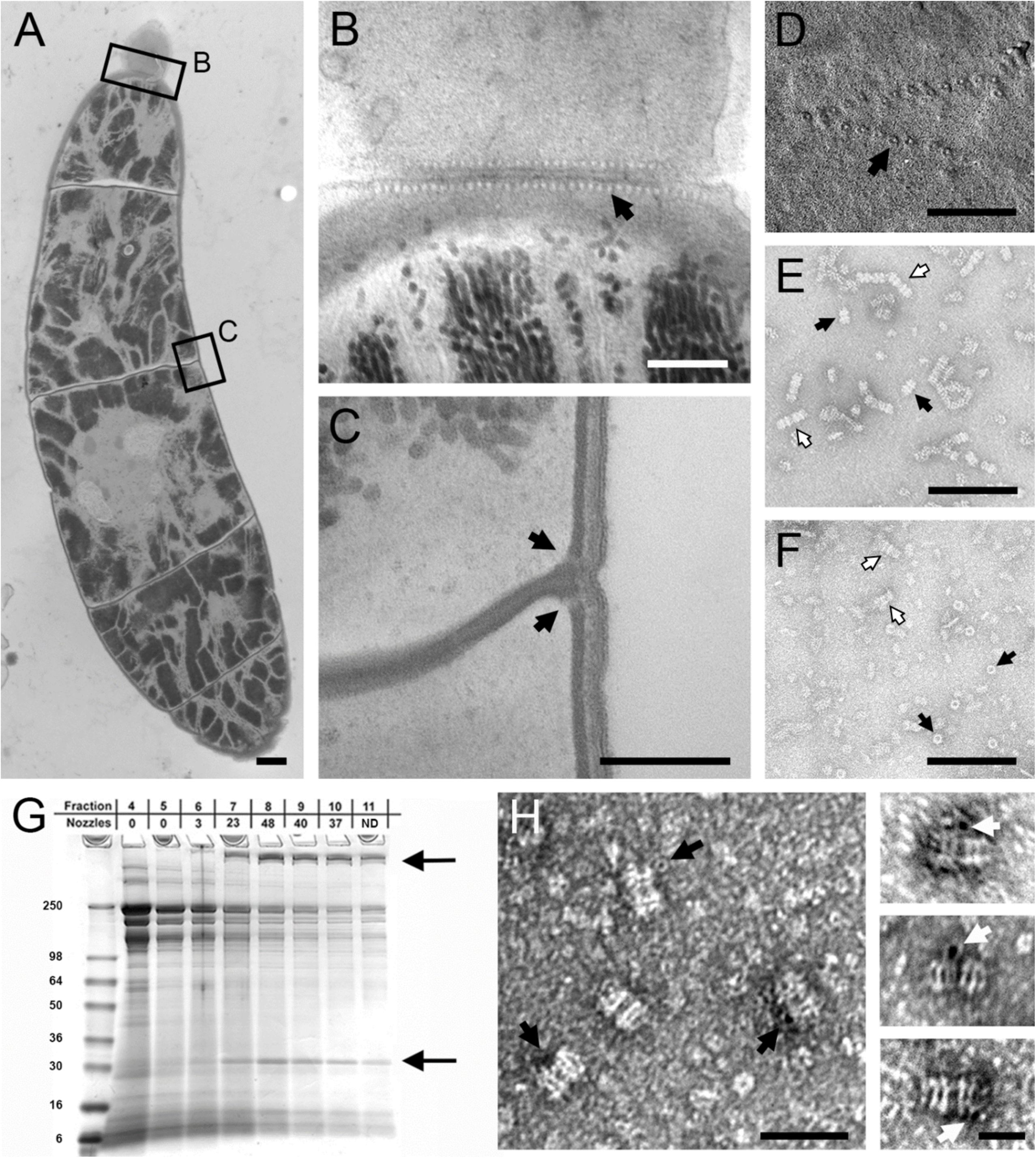
PilQ forms the slime nozzle in filamentous cyanobacteria (*A-C*) Thin sections of high-pressure frozen *Athrospira platensis* cells show the tilted transpeptidoglycan channels harboring the nozzle apparatus (black arrows), which are arranged in a circumferential ring at each cross wall. (*B*), (*C*) High magnification micrographs from indicated regions in (*A*). To visualize more channels, (*B*) shows the same region of the next thin section from a series of sections of the area shown in (*A*). The black arrows indicate the position of the transpeptidoglycan channels that in cross sections are visible as small white dots (*B*) and in longitudinal sections as slightly angled less dark stained bands traversing the peptidoglycan at an angle of 30-40° relative to the septum (*C*). (*D*) Pt/C shadowing of an isolated outer membrane patch reveals the rows of nozzles (black arrow) consisting of the peripheral ring (16-18 nm) and a central pore (6-8 nm) which have identical dimensions as the top views of the isolated pores (black arrows) in (*F*). (*E*) Transmission electron microscope image of a negatively stained isolated nozzle preparation. Black arrows indicate individual double ring nozzles, while white arrows indicate linear arrays containing multiple nozzles. The length of an individual double nozzle is ca. 32 nm. (*F*) As has been observed for nozzles of other filamentous cyanobacteria, adsorption to grids without glow discharge reveals top views of the complex (black arrows), while only a few side views are visible (white arrows) demonstrating that the cyanobacterial and myxobacterial nozzles (Fig. 3C) share similar architecture. (*G*) Fractions from a slime nozzle enrichment were screened by TEM and scored for how many nozzles were observed per grid square (ND, not determined). Fractions were separated by SDS-PAGE, and two bands correlated with fractions enriched for nozzles, identified as PilQ (black arrow higher mw band), and the pentapeptide repeat protein NIES39_A07680 (black arrow lower mw band). (*H*) Immunogold labeling of isolated PilQ nozzles showing overall labeling (black arrows) and binding of the 5 nm gold-labeled antibodies to individual nozzle complexes at higher magnification (white arrows in insets). Scale bars: (*A*) 1 μm; (*B*) 250 nm; (*C-F*) 200 nm; (*H*) 100 nm and 25 nm (insets).

With a candidate nozzle protein identified, we next attempted to visualize PilQ at the sites of slime secretion *in situ*. Although the ease of cultivation of *A. platensis* initially offered advantages, with continuous culture a substantial portion of the filaments lost their PilQ nozzles, ceased secreting slime, and became non-motile, a phenomenon that we had previously observed in permanently agitated cultures of benthic gliding cyanobacteria (62). As this mixed population of nozzle-containing and nozzle-free filaments yielded inconsistent results, we decided to use two highly motile benthic species, *Oscillatoria lutea* (SAG 1459-3) and *Phormidium autumnale* (strain Chesterfield) for further experiments. Genome sequence was obtained from both strains, and the gene for *pilQ* from *O. lutea* was expressed in *E. coli*. Protein was purified and used to inoculate rabbits to raise antisera. Although this antibody specifically cross-reacted with the PilQ band of both species in immunoblots (*SI Appendix*, Fig. S2*B*), initial attempts at fluorescent labeling of the nozzles in live filaments were unsuccessful. We attributed these difficulties to the inaccessibility of epitopes on PilQ due to the complex multilayered architecture of cyanobacterial cell envelopes. Here, the PilQ-containing outer membrane is sandwiched between a many nanometer-thick and heavily cross-linked peptidoglycan layer and an extracellular barrier comprised of an S-layer topped by the helically arranged glycoprotein oscillin (55, 62–63). To potentially increase access for the antibodies, we used isolated cell envelopes for our labeling experiments, but again failed to observe labeling of the PilQ nozzles at cell-cell junctions. However, within these preparations we consistently observed isolated disc-shaped cross walls that still had the nozzle-containing portion of the longitudinal wall attached (observed by the pores in the cell wall), and we saw clear peripheral immunolabeling of these cross walls with the anti-PilQ antisera (**Fig. 2*A***; *SI Appendix*, Fig. S4). These results supported our initial interpretation that PilQ epitopes were masked in our earlier attempts to immunolabel intact cells. Since a number of conventional permeabilization methods such as lysozyme or organic solvent treatment failed to allow labeling or resulted in the disintegration of the filaments, we attempted to perform limited cell lysis to remove some of the cell wall material. Exposure of live filaments to increased temperature or incubation with 200 mM DTT (64) were among the most reproducible treatments to induce limited cell lysis. Upon treatment of the filaments, the rows of nozzles were clearly labeled with the anti-PilQ antibody confirming that the nozzles at the cross walls were indeed composed of PilQ (**Fig. 2*B***). Unfortunately, the extensive multi-step treatment required for immunofluorescence imaging of the nozzles precluded the possibility to simultaneously retain and visualize slime secretion.

**Fig. 2.**
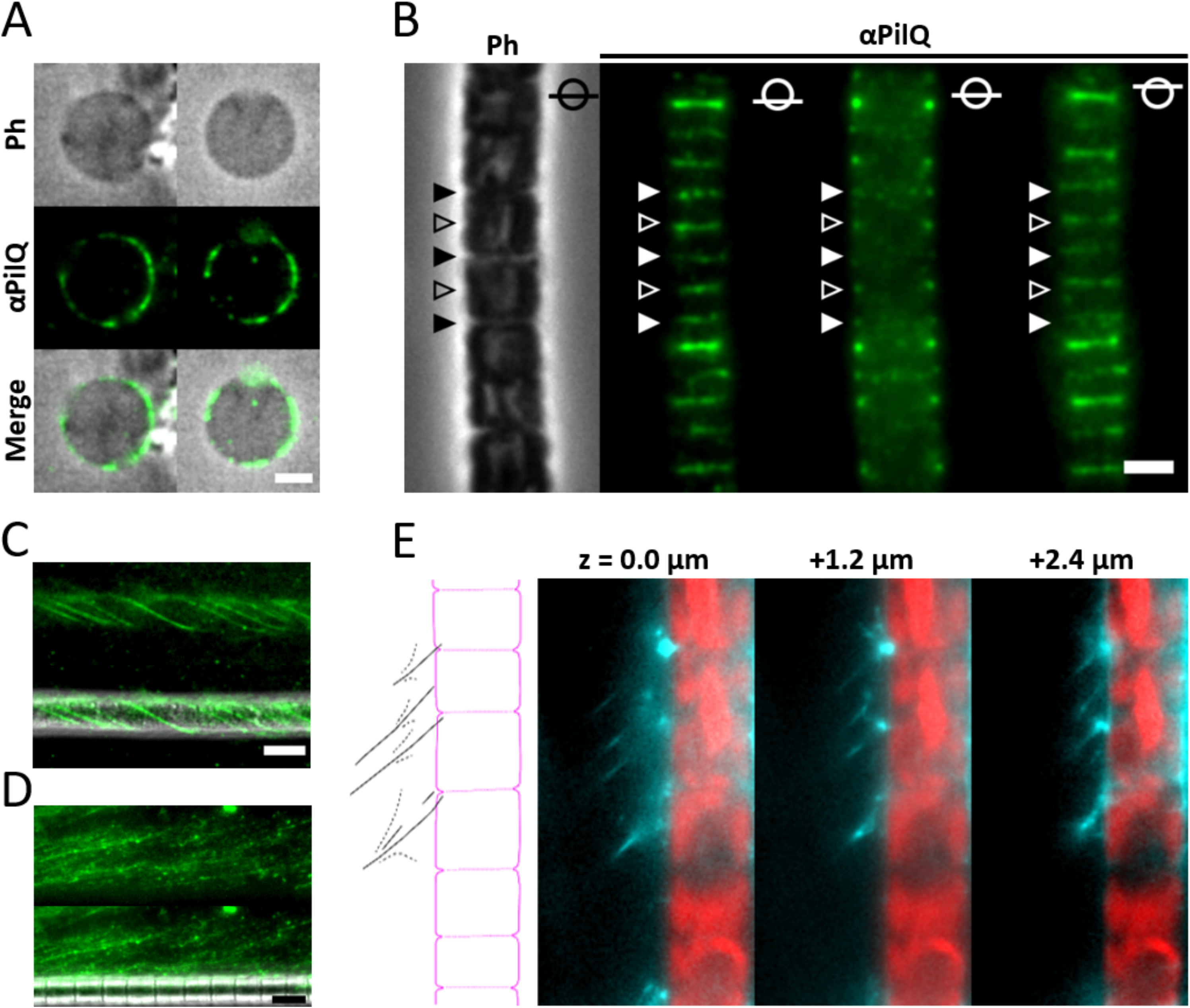
Immunofluorescence microscopy of *Oscillatoria lutea* and *Phormidium autumnale* filaments showing the localization of PilQ at the cross walls where EPS is secreted. (*A*) A PilQ antibody labels the periphery of isolated disc-shaped cross walls due to remnants of nozzle-containing longitudinal cell wall still being attached after cell breakage (*SI Appendix*, Fig S4). (*B*) Following limited autolysis, clear circumferential labelling is observed at the cross walls of the *O. lutea* filament. The image series are Z-stacks representing bottom, center, and top of the filament. Filled and hollow arrow heads denote mature and nascent cross walls, respectively. (*C*) Fluorescent concanavalin A labelling of live, motile *O. lutea* filaments typically showed strong adhesion of slime strands to filament surfaces. (*D*) Under strong lateral flow, strands of slime could be dislodged, but the location of their source is difficult to discern due to high gliding speed of cells and rate of slime secretion. (*E*) By subjecting *Ph. autumnale* filaments to brief sonication and cooling, individual strands of concanavalin A-labelled slime (green) could be observed to emanate from the cross walls where PilQ nozzles are located. Here auto-fluorescence at the center of the cell filament is displayed in magenta to more clearly show cell boundaries in this species. The schematic drawing on the left traces the outline of the cell filament (red) and the pattern of the lectin-labelled slime strands (cyan).

Consequently, we used fluorescently-labelled concanavalin A to visualize slime secretion in living cells to determine whether slime trails emerged from the cross walls, where nozzles are located. As the fluorescently labeled slime bands usually translocate along the filaments’ surfaces (**Fig. 2*C***), we had to apply a continuous flow to shear them from the surface. Under these conditions, the slime dislodged from the filament surface (51), revealing individual strands. However, the high gliding speed of these cells and the copious amount of slime secreted posed additional challenges in locating the precise origin of secretion (**Fig 2*D***). Subjecting the cell filaments to a gentle burst of sonication and cooling before imaging appeared to encourage slime dislodgement and decrease gliding speed, respectively. With these treatments, we observed individual strands of slime emanating in close proximity to mature and nascent cross walls (**Fig, 2*E***), where PilQ nozzles are located (**Fig. 2 *A* and *B***). This is consistent with a previous report of the localization of slime secretion when slime was stained using India Ink (51). Taken together, this evidence supports the interpretation that the secretin PilQ is used for slime secretion in filamentous cyanobacteria.

### GspD is a Candidate for Slime Nozzles in *M. xanthus*

Because multicellular filamentous cyanobacteria are difficult to genetically manipulate, and to test if other slime-secreting bacteria also use secretin nozzles, we next studied the soil bacterium *M. xanthus*. This strategy was based on earlier observations of virtually identical nozzle-like structures in the outer membrane of this bacterium that were in close proximity to the emergence of slime bands on the surface of the cells (52). To identify the nozzles from *M. xanthus*, we used a similar approach as for the cyanobacteria. Isolated cell envelopes were purified and solubilized. We examined fractions by electron microscopy to screen for the presence of structures of similar shape and size as the OM-embedded nozzles previously observed (**Fig. 3*A***). In contrast to the nozzles from *A. platensis*, we only observed ring-shaped top views of the complex, but not side-views (compare **Fig. 1*F*** and **3*C***). Using our fractionation protocol, we isolated fractions highly enriched in nozzle-like structures, and correlated the presence of these nozzles to a ∼270 kDa band on SDS-PAGE gels (**Fig. 3 *B* and *C***). Using mass spectrometry, the band was identified as GspD (*SI Appendix,* Table S1), suggesting that secretins are also used by myxobacteria in slime secretion. Since the secretin PilQ in *M. xanthus* is known to contribute to social (S-) motility as the outer membrane channel of the type IV pilus (65–67), but not gliding motility, we tested whether PilQ was also used for slime secretion in this species. Using mutants that lack PilQ, we successfully isolated nozzles and observed slime trails that were indistinguishable from the wildtype, demonstrating that PilQ is not involved in slime secretion (*SI Appendix*, Fig. S5). Of note, like in the investigated cyanobacteria (**Fig. 1*G*** and *SI Appendix*, Fig. S2), the molecular weight of the PilQ/GspD band was substantially larger than the predicted molecular weight of the mature outer membrane-associated protein (i.e. *A. platensis*: 756 aa, 81 kD; *M. xanthus*: 840 aa, 90 kDa). Moreover, the high-molecular-weight bands from both species displayed a pronounced temperature-dependency; while the intensity of the *A. platensis* PilQ band decreased somewhat upon boiling, the *M. xanthus* GspD band completely disappeared after heating above 70 °C. We interpret these observations to indicate that at high concentrations and high temperatures, the protein irreversibly aggregated and failed to enter the gel (68). By contrast when using smaller amounts of GspD that are present in whole cell lysates and visualized by immunoblot, neither the high-molecular-weight proteins nor the temperature-dependence were observed (compare **Fig. 3*B*** and **4*A***). Under these circumstances, we observed a protein band at the expected molecular weight of ∼100 kDa.

**Fig. 3.**
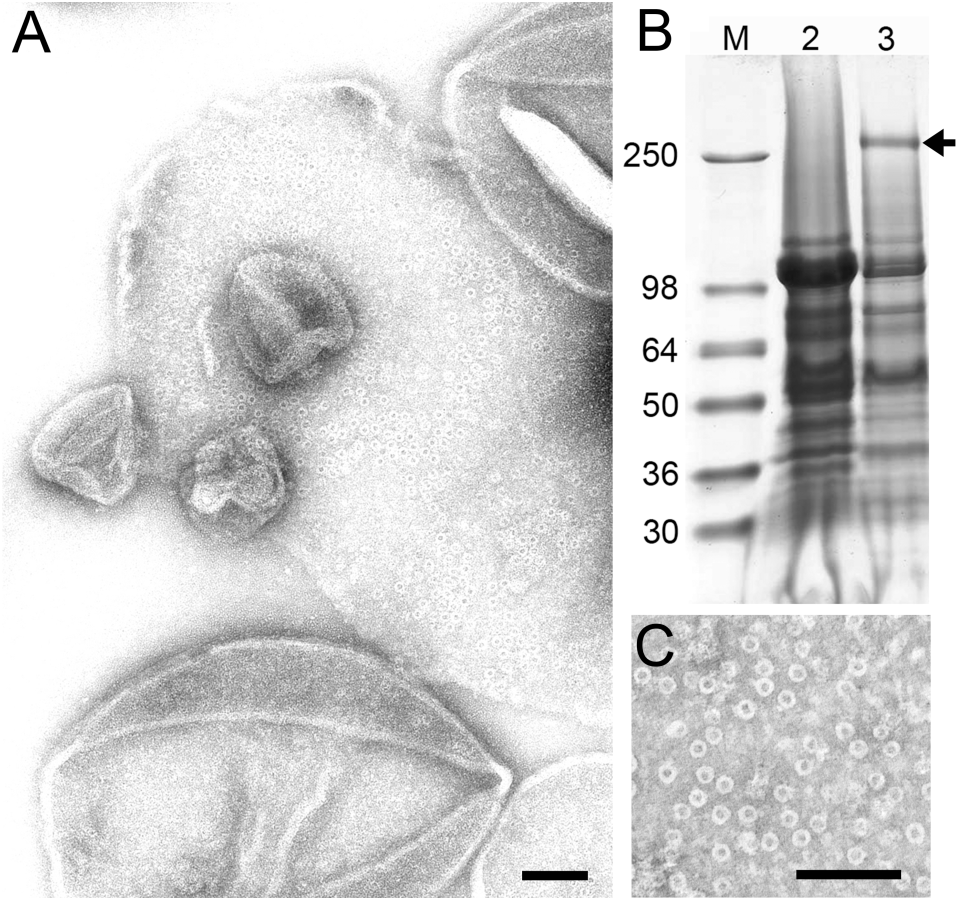
Isolation and identification of GspD in *Myxococcus xanthus* (*A*) Electron micrograph of nozzle-like structures *in situ* in outer membrane fragments from *M. xanthus* at high magnification. (*B*) SDS-PAGE of the purified nozzle preparation showing the ∼270 kDa band that is formed after heating highly concentrated samples of GspD at 70 °C. Lane 2, biochemical fraction lacking nozzles (as evaluated by electron microscopy). Lane 3, biochemical fraction containing nozzles. Arrow indicates protein band unique to fractions enriched for nozzle-like structures, and identified as GspD *SI Appendix*, Table 1). No other protein was consistently found to co-purify with nozzle-like structures in these isolations. (*C*) Electron micrograph of a purified isolation of the 14-16 nm wide ring-shaped nozzle complexes formed by GspD. Scale bars: (*A*, *C*) 100 nm.

**Fig. 4.**
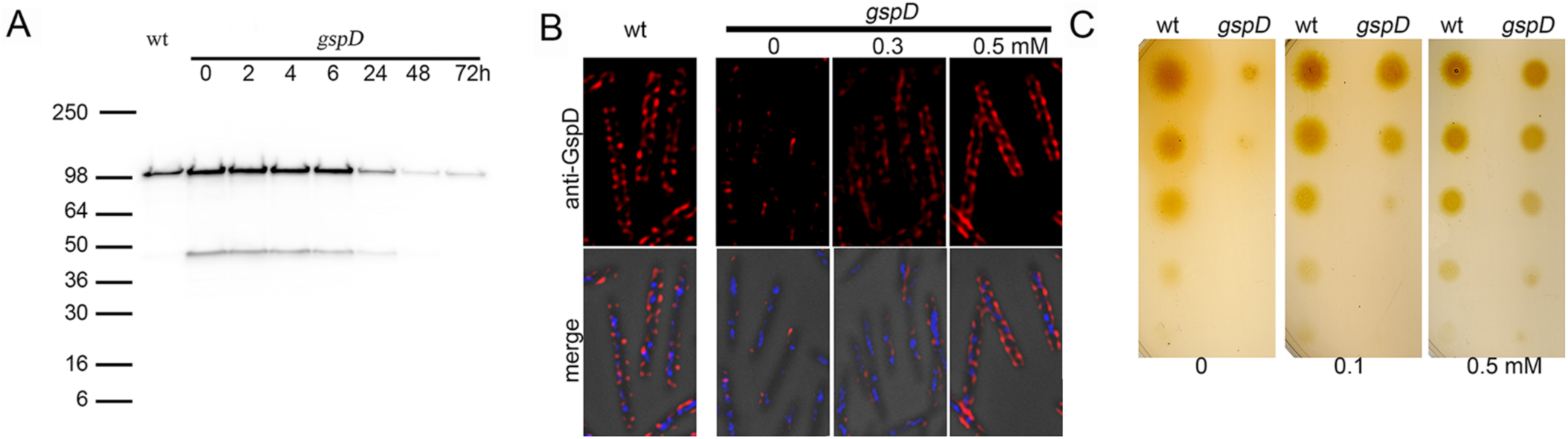
*gspD* is an essential gene in *Myxococcus xanthus* (*A*) Immunoblotting reveals that removal of copper from the medium of a cell line expressing *gspD* under a copper-inducible promoter leads to a gradual decrease of GspD, which eventually stabilizes at a low but detectable level. GspD is visualized as a ∼100 kDa band, and occasionally a second band at ∼47 kDa. (*B*) Immunofluorescence microscopy shows that GspD signal is due to low-level expression of the protein in all cells. Cells were grown in the absence or presence of 0.3 mM CuSO_4_ for 24 h prior to fixation and imaging. Top panel, anti-GspD signal; bottom, merged image with anti-GspD (red), DAPI (blue), and phase contrast. (*C*) Plating of wild type cells and cells containing a copper-inducible version of GspD reveal that *gspD* is an essential gene. Cells pre-cultured in 0.2 mM CuSO_4_ were shifted to culture lacking CuSO_4_ for 48 h. Cells were then concentrated, and 4-fold serial dilutions were spotted on agar media containing the indicated concentration of CuSO_4._ In the absence of copper (left panel) the mutant strain fails to grow, while with increasing amounts of copper (intermediate concentration, middle panel; high concentrations, right panel) the cells grow at rates indistinguishable from the wild type.

### *gspD* is an Essential Gene in *M. xanthus*

To study a possible contribution of GspD to *M. xanthus* slime secretion, we attempted to generate a markerless deletion mutant of *gspD*. However, while we were able to recover multiple clones with an integrated deletion plasmid, we consistently failed to recover a deletion mutant following a second recombination event to remove the plasmid. Instead, all attempts yielded clones that had reverted to the parental wildtype strain, a result we obtained across multiple attempts in different genetic backgrounds. We next pursued a strategy of generating a conditional knockdown mutant. For this, we introduced a second copy of *gspD* under the control of the copper-inducible promoter, P*_cuoA_* at the *attB* site (69) into our clones that had successfully integrated the deletion plasmid. When selecting for removal of the plasmid in the presence of copper, we were able to recover multiple clones with *gspD* deleted from the chromosomal locus. These observations support the interpretation that *gspD* is an essential gene.

To test the depletion of GspD, we grew cultures in media with copper, then washed and re-suspended the cells in media lacking copper, but containing the copper chelator bathocuproinedisulfonic acid (BCS). Equal cell numbers were collected at various time points, lysed with sample buffer, and examined by immunoblot using an affinity purified antibody against the C-terminus of GspD (see materials and methods for details). GspD levels declined for more than 24 h following removal of copper before leveling off at a low, but consistently detectable, amount (**Fig. 4*A***). This low level was not due to a small number of escape mutants, but was visualized by immunofluorescence as a weak signal in all cells present in the culture (**Fig. 4*B***). Cells grown in the presence of high concentrations of copper displayed enhanced fluorescence at the periphery of the cell, in a pattern consistent with signal from endogenous GspD in wildtype cells but at levels higher than for endogenous protein (**Fig. 4*B***). Overexpression of GspD under these conditions was similarly confirmed by immunoblot (*SI Appendix*, Fig. S6).

Consistent with the expression patterns of GspD in copper-depleted cells, we found that our *gspD* cells would grow for several generations in liquid culture in the absence of copper, but at longer times (>24 h) the growth rates of the cultures would dramatically decline. To test the requirement for copper in the media, cells were grown in the absence of copper for 48 h (the earliest observed time of maximum GspD depletion (**Fig. 4*A***)), and serial dilutions were spotted on agar plates lacking or containing copper. We observed no effect of this handling on the survival or growth of wildtype cells, but *gspD* mutants were highly dependent on copper in the media, confirming that the cells need to express GspD in order to survive and grow (**Fig. 4*C***).

### GspD Depletion Yields Fewer Nozzles and Reduced Slime Secretion in *M. xanthus*

To test the hypothesis that GspD is the major component of the slime nozzle, we grew *gspD* mutant cells in the absence or presence of copper. Cells were collected, and OMs were disrupted with glass beads and examined by TEM for the presence of nozzles (52). While we found few of the complexes in the OM from cells depleted for GspD, we observed large numbers of such structures in the OM fragments from cells grown in the presence of copper (**Fig. 5*A***).

**Fig. 5.**
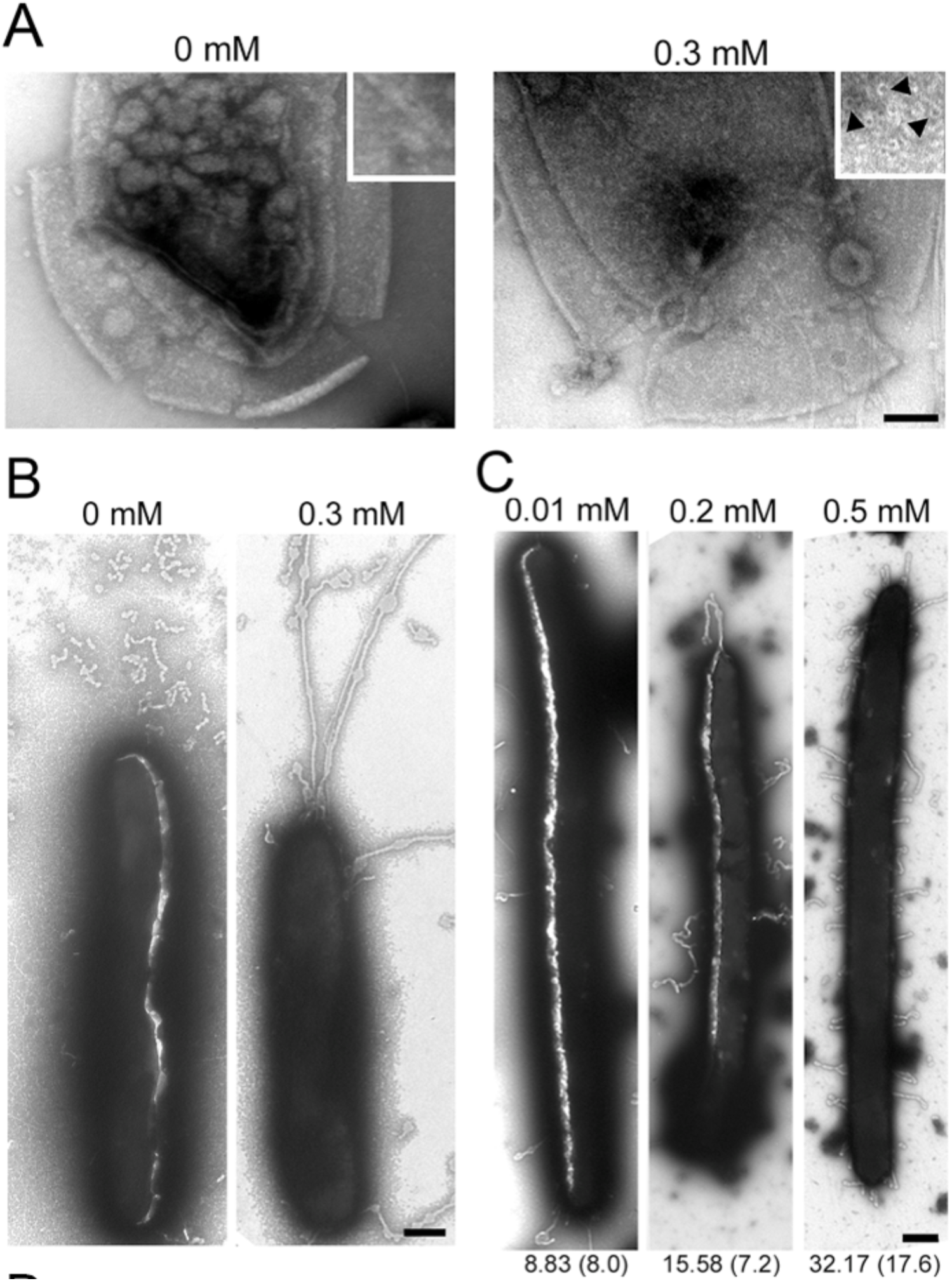
Depletion of *gspD* results in a reduction of the number of nozzles (*A*) Electron micrograph of negatively stained outer membrane fragments from disrupted cells, revealing few or numerous ring-shaped GspD complexes depending on the absence or presence of copper. The insets show representative areas of the outer membranes at higher magnification. Black arrows indicate GspD complexes. Scale bar, 100 nm. (*B*) Electron micrograph of representative cells in the absence or presence of copper. Cells that have very few GspD nozzles secrete little or no slime, while cells possessing normal numbers of GspD nozzles secrete easily detectable bands of slime. (*C*) Cells grown in 0.2 mM CuSO_4_ were shifted to the indicated concentration of CuSO_4_ for 24 h. Cells were allowed to swarm on grids, and visualized by electron microscopy. For each condition, at least 12 cells were scored for the number of slime trails emanating from the bodies of isolated cells with intact membranes. The mean number of slime trails per cell ± S.D. is presented. All concentrations are mM CuSO_4_. Scale bar, 500 nm.

We next wished to assay for production of slime. Multiple methods have been reported for the detection of slime in *M. xanthus*, including phase contrast microscopy (70), India ink (71), acridine orange (52), atomic force microscopy (72), wet-SEEC or fluorescently labeled ConA (54). However, material other than slime is produced by cells during locomotion and biofilm formation (73–75), which may confound results. Thus, to visualize slime directly, we performed negative staining and examination by electron microscopy, as described (52). To observe slime trails, we grew cells in liquid culture with or without copper for 40 h, spotted them on EM grids coated with hydrolyzed chitosan, and allowed them to glide. Grids were then stained and examined by TEM for the presence of slime trails. We identified slime trails as having distinct morphology (distinguishable from membrane vesicles and tubule-like outer membrane protrusions, as well as the S-motility-related fibrils) in the TEM, and by their pH sensitivity, as treatment with acidic stains (un-buffered uranyl acetate (UA), pH 4.5 or SiPTA at pH 4.0) removed slime trails (but not other membrane components, i.e. vesicles) from grids, while neutral stains (SiPTA at pH 7.0) did not (*SI Appendix*, Fig. S7). We consistently found that cells expressing *gspD* regularly secreted slime, visualized as persistent and thick trails emerging from the cell body, whereas cells depleted for GspD produced very low levels of slime, or none at all (**Fig. 5*B***). In these depleted cells, the only extracellular material that resembled slime was often fragmented bands of material, thinner and shorter than slime trails observed in wildtype or *gspD*-overexpressing cells (**Fig. 5*B***).

We considered that the loss of the essential functions of GspD may lead to cell death, and the lack of slime secretion we observed was simply due to observations of dead or dying cells. To address this concern, we grew cells in the presence of low, moderate, or high concentrations of copper for 24 h. We selected 24 h as the time for pre-culture, since at this time point, there is depletion of GspD from the cells, but not maximal depletion (**Fig. 4*A***), and cells in liquid cultures did not yet show a growth defect. We selected copper concentrations that had previously demonstrated minimal toxicity to *M. xanthus* cells (69). Cells grown under these conditions were spotted on EM grids, and the numbers of slime trails emerging from individual cells with intact membranes (to avoid sick or dead cells) were counted. Compared to the cells grown with moderate copper levels, cells grown with low levels of copper produced nearly half as many slime trails (15.6 ± 7.2 *vs*. 8.8 ± 8.0 trails/cell). Moreover, overexpression of GspD resulted in a doubling of the slime trails for cells grown in high levels of copper (32.2 ± 17.6 trails/cell; **Fig. 5*C***). This observed GspD-dependent increase is a strong indication of the direct contribution of GspD to slime secretion.

Taken together, these data demonstrate that under conditions where GspD was partially depleted from the cells, that cell survival was unaffected, whereas slime secretion was dramatically reduced. These data support the conclusion that slime secretion is specifically associated with the reduction of GspD.

### GspD is Necessary for Gliding Motility in *M. xanthus*

As all models for gliding motility in *M. xanthus* suggest an important role for slime (53, 76), we predicted that slime-deficient mutants should be defective in gliding. To test this, we grew cells for 24 h in media lacking copper, spotted these cells onto agar plates containing, or lacking copper, and allowed cells to swarm for 48 h. When these cells were plated onto media lacking copper, we observed cell growth from the initial, dilute spot. However, while the absence or presence of copper had no effect on the ability of wildtype cells to expand, the *gspD* mutant completely depended on copper for individual-cell motility (**Fig. 6**). To ensure that *gspD* expression was stimulating gliding (adventurous, or A-motility in *M. xanthus*), and not the type-4 pilus-dependent S-motility, we generated *gspD* mutants in the S-motility deficient Δ*pilA* background. Whereas the parent strain was able to expand in the absence or presence of copper, the *gspD* mutant required copper for motility (**Fig. 6**).

**Fig. 6.**
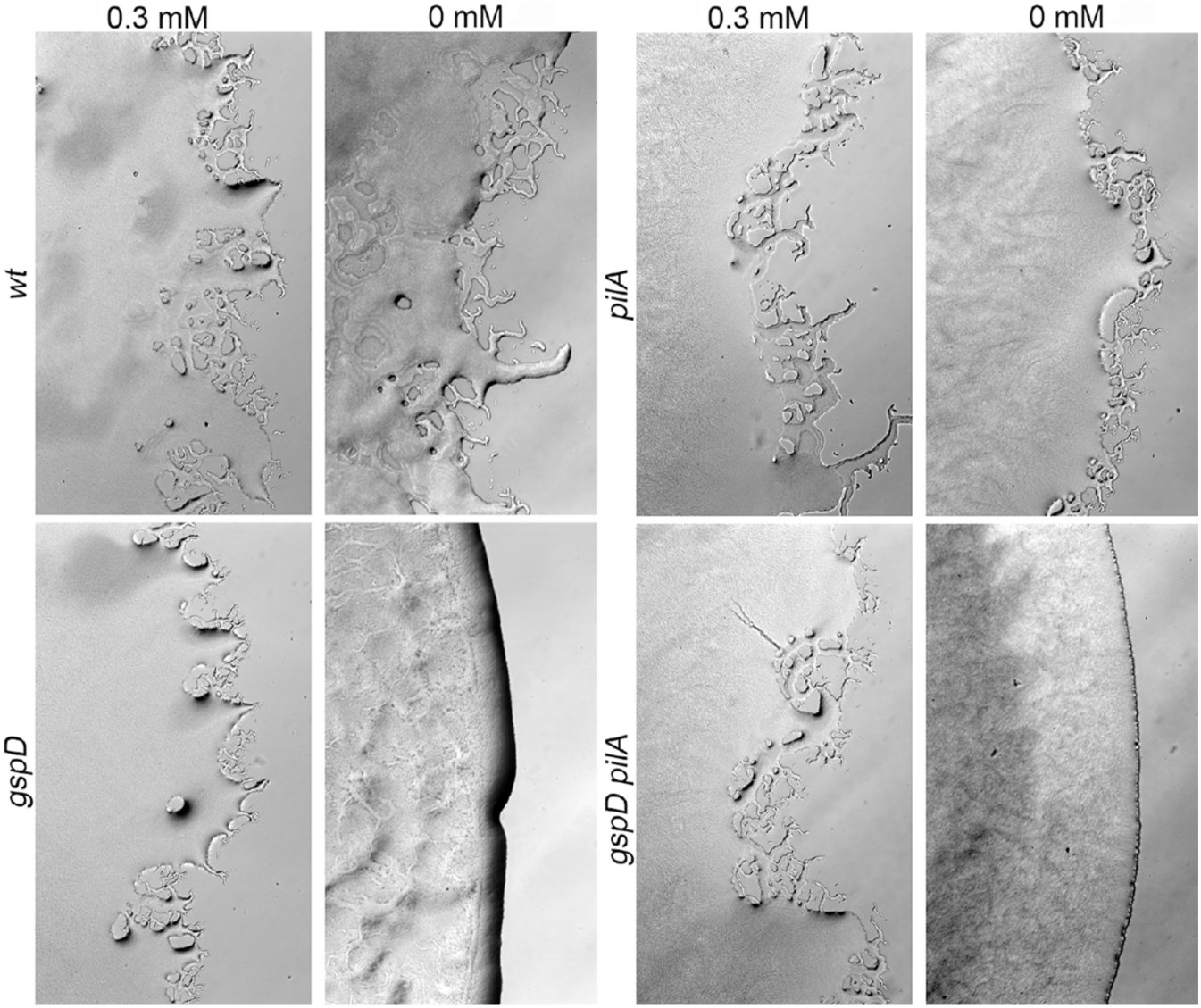
GspD is required for adventurous (A-) motility in *Myxococcus xanthus* Appearance of the swarm colonies and edges of wild type, *gspD* cells, Δ*pilA*, and *gspD* Δ*pilA* cells in the presence and absence of copper. Single cell-based A-motility, characterized by individual cells spreading from the colony edge outwards, can be observed in the wild type cells and the Δ*pilA* cells, while the depletion of GspD abolishes this movement in the *gspD* and the *gspD ΔpilA* cells, confirming that GspD is required for A-motility.

Since we had concluded that *gspD* is an essential gene, we tested that in this assay the cells were living, but simply unable to glide. All of the swarm colonies became denser over the 48 h of the assay, including those that did not demonstrate motility, indicating growth of the colonies. Moreover, *gspD* cells grown under conditions similar to the gliding motility assay, but plated on soft agar to promote S-motility, demonstrated swarm expansion typical of S-motility in both the absence and presence of copper (*SI Appendix*, Figure S8*A*), demonstrating both that GspD was not necessary for S-motility and that even under conditions of GspD depletion, cells were still actively motile. The swarm colony was smaller for cells grown in the absence of copper, likely due to a slower growth rate of the cells from depleted levels of GspD; however, motility was clearly observed. We also collected cells from Δ*pilA gspD* swarm colonies plated in the absence or presence of copper, and determined cell viability. We observed no differences in the ratio of living to dead cells (*SI Appendix*, Fig. S8*B*), suggesting that the cells that survived the copper depletion expressed enough GspD to survive, but not to swarm (**Fig. 4*A***). Taken together, these results demonstrate that cells sufficiently survived the depletion of GspD in these experiments, and that swarm expansion could have been detected had it occurred. Thus, we conclude that *gspD* is an A-motility gene, which may have not been identified in previous genome-wide genetic screens (reviewed in (77)) because it is an essential gene.

## Discussion

EPS secretion is an important strategy for environmental adaption of bacteria (78). With enormous varieties of chemical compositions, molecular weight, and adherence to bacterial surfaces, these molecules serve a wide variety of purposes, including as important components of the bacterial cell envelope (4, 26, 32), providing protection against desiccation and toxic substances (78–80), mediating attachment to surfaces (78, 81), biofilm formation (82–83), host interaction (84–85), and bacterial motility (53, 56). Although there are many methods for detection of bacterial EPS, relatively little is known about the chemical composition, synthesis, and secretion of these molecules.

Here we show that the secretins PilQ and GspD form the previously observed EPS-secreting nozzles in cyanobacteria (51) and myxobacteria (52). As Gram-negative bacteria can possess multiple envelope-associated macromolecular secretory complexes, it was essential to ensure that the ring-shaped molecules we isolated were indeed the slime nozzles. For this reason, we initially used the cyanobacterium *A. platensis*, which, as a photosynthetic autotroph, is capable of EPS production while having fewer secretory systems that may have been mistaken for nozzles. In fact, BLAST searches reveal that none of the three cyanobacteria species used contain transport systems with large outer membrane gate structures such as T3SS, T4SS, and T6SS. Only Wza homologs are found that are substantially smaller than the nozzles, based on the dimensions of the *E. coli* protein (outer diameter 4.6 nm). In line with these observations, isolations from *A. platensis*, *O. lutea*, and *Ph. autumnale* invariably yielded a single type of ring-shaped complex formed by PilQ, allowing identification of secretins as the principle structural component of the slime nozzles of filamentous cyanobacteria (51). This interpretation is supported by immunoblot analyses (*SI Appendix*, Fig S2), mass spectrometry (*SI Appendix*, Table S1), structural comparisons with known secretin complexes (*SI Appendix*, Fig S3; **10**), direct immunogold labeling of the isolated complexes (**Fig. 1*H***), immunofluorescence microscopy of *O. lutea* and *Ph. autumnale* filaments (**Fig. 2 *A* and *B***), and the correlation of the localization of PilQ (by immunolabelling) with the pores (by electron microscopy) to isolated cross walls (compare **Fig. 2*A*** and *SI Appendix*,Fig S4).

The identification of the secretins PilQ and GspD as the OM channels for EPS secretion in cyanobacteria and *M. xanthus*, respectively, was consistent with the electron microscopic appearance of the isolated nozzles (10). In contrast, the presence of the second protein in cyanobacteria, the pentapeptide repeat protein NIES39_A07680 was surprising. Pentapeptide repeat proteins form a family (Pfam 00805) whose members are not widely distributed beyond cyanobacteria and have been implied in unknown targeting or structural functions (86). Although NIES39_A07680 appeared to co-purify with PilQ, its structural or functional relation to the secretin is currently unknown. Its small size and the presence of multiple repetitive putative protein-protein interaction motifs (pentapeptide repeats (86) rather than TPR (87)) reveal that NIES39_A07680 shares no similarity with known pilotins or secretin accessory proteins (20). Another unexpected finding was that the nozzles we recovered from cyanobacteria were almost entirely dimers of PilQ rings (615 out of 618 structures were dimers), similar to earlier observations (333 out of 334 structures; **51**), while the *M. xanthus* nozzles were isolated as single ring complexes. Although initially quite different in appearance (compare **Fig. 1*E*** and **3*C***), adsorption of *A. platensis* nozzles to grids without glow discharge resulted in top views that clearly revealed the common ring-shaped architecture of the nozzles (compare **Fig. 1*F*** and **3*C***). We considered two scenarios that could account for the different appearance of the cyanobacterial nozzles: the cell envelope-embedded structures may in fact be monomeric rings as in all other studied secretins and the observed dimers had formed during their isolation, or the nozzles are dimers, revealing plasticity in certain secretins to form novel structural arrangements. The isolation of nozzles from *M. xanthus* may have been confounded by the presence of additional OM translocation machineries. However, our isolations fortuitously contained only one type of ring-shaped complex, namely the secretin GspD. In contrast to the nozzles from cyanobacteria, nozzles from myxobacteria were exclusively recovered as single ring structures, and no proteins were co-purified. These observations support the interpretation that the nozzles in both cyanobacteria and *M. xanthus* are monomeric rings that lack associated proteins or pilotins when fully assembled, and the visualization of PilQ dimers and recovery of NIES39_A07680 were likely artifacts of the sample preparation.

As PilQ is a component of the type II secretion system of the type IV pilus apparatus, our findings were consistent with the discovery that type IV pili-related proteins localize to the cross wall of the *Nostoc punctiforme* hormogonia and contribute to their transient motility (88). From our own and published genomic analyses, cyanobacteria belonging to the order Oscillatoriales, like the three species in this study, contain all components of the type IV pilus motility machinery required for function, and only one copy of *pilQ* is found. However, we have not observed pilus or pilus-like surface appendages in Oscillatoriales in this or previous studies (51, 55, 62). Although our findings appear to contradict earlier descriptions of putative pili (“fimbriae”) in *Arthrospira* and *Oscillatoria* (89), the pili described in this earlier study fundamentally differed from all other known unicellular cyanobacterial and prokaryotic pili: they were described as a helically arranged, tightly attached array consisting of parallel filament-like elements covering the entire surface of these filamentous species (89). While some investigators interpreted the parallel running elements of the array as pili (89), others thought of them as contractile actin-like filaments involved in the gliding motility of *Oscillatoria* species (90). Of note, the characteristics of these surface arrays, their tight attachment to the surface, the parallel arrangement of their substructures, the 60° angle with which the individual elements run helically along the long axis and the diameter of the individual substructures (6-9 nm) (89, 90) are identical to the features of the extracellular surface layer formed by the glycoprotein oscillin in *Oscillatoria*, *Lyngbya*, and *Phormidium* species (62). Importantly, oscillin is a large (> 66 kDa in *Ph. uncinatum*), heavily glycosylated, Ca^2+^-binding protein mostly composed of *β*-sheets that does not share any similarity with the small (< 25 kDa) *α*-helix-containing pilins. Taken together, we conclude that the structures described in this study are most likely the oscillin array, and more evidence would be needed to establish the presence of pili in members of the Oscillatoriales. Nonetheless it may be possible that, like *M. xanthus*, some filamentous cyanobacteria like *Nostoc* can switch between pilus-dependent and -independent modes of motility, but under yet unknown circumstances, which intriguingly suggests that secretins represent a conserved core component that is important to both gliding and pilus-dependent motility.

Unlike the three studied cyanobacteria, the predatory myxobacteria (91–92) possess copies of virtually all known Gram-negative protein secretion machineries (93). In fact, the genome of *M. xanthus* contains 3 paralogs of *gspD*, namely *gspD*, *pilQ*, and *mxan_RS15055* (previously *mxan_3106*) (93), with the greatest sequence similarity within the secretin domain (*SI Appendix*, Fig. S9). All three paralogs have been identified in proteomics studies as expressed and localized to the OM or OM vesicles of *M. xanthus* cells under various environmental conditions (94). PilQ is the outer membrane secretin of the type IV pilus (66), while Mxan_RS15055 (Om031 in *M. fulvus*) has been reported to be involved in osmoregulation allowing cells to better survive under increasing salinity (95). Together, these data show that the three paralog secretins, PilQ, Mxan_RS15055, and GspD have distinct non-interchangeable functions and that the role of GspD in slime secretion and A-motility is unique.

To explain the dependence of slime secretion on the presence of secretins, we consider two plausible mechanism: secretins could either be directly involved in slime secretion as the OM gates through which the synthesized polymer is secreted, or indirectly by secreting enzymes that then polymerize slime on the cell surface, similar to synthesis of bacterial dextranes (28). In dextrane production, secreted surface-associated transglycosylases enzymatically cleave extracellular sugar polymers such as sucrose, starch, or fructanes to convert the resulting monosaccharides into dextran polymers. To consider whether such a process could account for slime polymerization in our organisms, it is important to study the repertoire of secreted proteins. In *M. xanthus*, GspD has recently been shown to translocate MYXO-CTERM domain-containing proteins (96), of which 34 have been bioinformatically identified using the TIGR03901 consensus motif (97). Only one of those 34 proteins, MtsC (Mxan_RS06455, MXAN_1334; **98**) is involved in motility, but not A-motility. None of the five MYXO-CTERM domain-containing proteins, (Mxan_RS04600, MXAN_6274, PQQ-dependent sugar dehydrogenase; Mxan_RS30220, MXAN_6236, putative polysaccharide-degrading enzyme; Mxan_RS30405, MXAN_6274, polysaccharide deacetylase family protein; Mxan_RS34095, MXAN_7044, exo-alpha-sialidase; Mxan_RS34570, MXAN_7140, glycosyl hydrolase) that are involved in carbohydrate metabolism show similarity to transglycosylases. Moreover, CTT does not contain cleavable sugar polymers, and physiological experiments have shown that *M. xanthus* is unable to utilize glucose, starch, or glycogen from the medium (99). Together, these observations indicate that is highly unlikely that the slime in *M. xanthus* could be synthesized using an extracellular transglycosylase reaction (28). Likewise, in filamentous cyanobacteria, no transglycosylases (or indeed, any proteins) have been identified as PilQ substrates that could polymerize slime outside of the cell. BG11 medium, like CTT, does not contain any carbohydrates that could act as substrates for transglycosylase-like enzymes. To allow extracellular polymerization in the absence of cleavable carbohydrate precursors would necessitate the secretion of large quantities of activated UDP-sugars by the bacteria. However, no such polymerization process has been reported in any bacterium, and the unavoidable loss of UDP would make such a process metabolically extremely costly. Therefore, we consider the most plausible interpretation of our findings to be that the role of the secretins is to secrete polymeric slime. The discovery that bacteria use secretins as the OM gate for EPS secretion prompts the question whether this mechanism represents a completely novel type of EPS secretory pathway, or whether the secretin is used as the OM component of other, already known, EPS secretions systems (7, 28). Our attempts to test if additional components of the Gsp machinery contributed to slime secretion in *M. xanthus* (GspE, GspG, and GspK) failed, as we were unable to obtain markerless deletions in the corresponding genes, suggesting that these are all essential genes, similar to *gspD* (own, unpublished observations). We also do not yet know all proteins involved in the synthesis, polymerization, and trans-periplasmic transport of slime. Nonetheless, evidence from cyanobacteria indicates that slime secretion may involve a synthesis-dependent mechanism. The secreted slime in *Phormidium uncinatum* is a complex heteropolysaccharide (100), and recent genetic work has identified a highly conserved, 13 gene-long locus that is important for EPS secretion and motility in all sequenced filamentous cyanobacteria (101). This *hps* locus encodes for five glycosyl-transferases (*hpsEFG*, *I*, and *K*) and four pseudopilins (*hpsBCD* and *H*), among others. The involvement of these genes suggests a potential link to secretins, as pseudopili have been proposed to act as pistons to push protein cargos through the OM secretin gate (22).

Therefore, it is tempting to speculate that the *hps* locus encodes parts of a novel synthase-dependent system that secretes EPS slime using PilQ/GspD as the OM gates. Of note, the pore size of the secretin gates is substantially larger (6-8 nm) than the opening of other carbohydrate secretion gates, such as the Wza channel (1.7 nm (38)) or the alginate secretion porin AlgE (0.8 nm (102)). As alginate, for example, is a high-molecular-weight carbohydrate, the pore diameter does not appear to correlate with the molecular weight of the secreted EPS, suggesting that other factors dictate the size of the OM gate for a given secretory system. One such factor may be the number of polymer strands that are simultaneously secreted through the channel, suggesting that secretins may be high-throughput gates allowing the rapid secretion of multiple strands of EPS that could form the electron microscopically observed ribbons (this work and **52**).

The fact that GspD in *M. xanthus* is involved in two very different secretory processes, namely slime and protein secretion, may explain why genome-wide genetic screens have not identified mutant strains that were completely deficient in slime secretion, indicating that one or both of these processes are essential (70, 77). If slime secretion is necessary to the cell (for example, in formation of a capsule) or secretion of the MYXO-CTERM domain-containing proteins is essential (as they are essential surface-associated proteins (96)), this would explain our observation that *gspD* is an essential gene.

Intriguingly, this raises the possibility that the same OM channel might engage multiple “accessory” protein complexes in the periplasm and cytoplasmic membrane. This potential versatility may explain why there appears to be a mismatch between the number of GspD nozzles and their distribution across the cell body with the observed slime bands emerging from the cell surface (an average cell possesses about 250 nozzles per pole (52) and a somewhat lower number spread over the length of the cell, while we observed many fewer slime bands; see i.e. **Fig. 4*B*** and *SI Appendix*, Fig. S5). A substantial number of GspD secretins may therefore participate in protein secretion alone, or multiple nozzles may contribute to each slime band. While plausible, this scenario is not the only possible explanation. It may be that slime secretion itself is essential in order to balance metabolic fluxes, or that GspD is involved in so far unknown transport process such as the release or uptake of low-molecular-weight substances, a possibility that is supported by observations of the diffusion of small molecular weight substrates through “closed” secretins (25, 103).

An important aspect of EPS secretion in cyano- and myxobacteria is its putative role in gliding motility in these organisms (52, 56). Although an important role for slime secretion for motility is generally accepted (53), its exact contribution is a matter of debate ranging from a passive adhesion factor (54), to a viscoelastic substrate (76), to a propulsive force generator (51, 101). What complicates resolving these issues is the possibility that the contribution of slime secretion to motility may be different in different bacteria. For the normally non-motile cyanobacterium *N. punctiforme*, hormogonia (short, transiently motile filaments) were recently reported to use slime secretion and type IV pilus-related proteins in gliding motility (88). Based on the finding that mutant strains lacking multiple glycosyltransferases (HpsE-G) were deficient in motility, and the observation that media conditioned by wild-type hormogonia could restore motility in these mutants, it was suggested that slime secretion facilitates motility but does not generate the motive force for gliding in *N. punctiforme* hormogonia (88). Importantly, permanently motile filamentous species like *O. lutea* and *Ph. autumnale*, or the previously studied cyanobacteria of the genera *Oscillatoria*, *Phormidium*, *Lyngbya,* and *Anabaena*, lack type IV pili (51, 55, 89) but still possess the conserved *hps* locus (101). We suggest that these species may synthesize slime similar to *N. punctiforme*, but may use their secretin PilQ directly for its secretion. Moreover, the absence of pili precludes that either retraction (like in *Synechocystis* or *Myxococcus*) or extension (as suggested for *N. punctifome*) of these structures could power movement in the vast majority of filamentous cyanobacteria that, like the aforementioned cyanobacteria, are permanently motile but without pili. An important unresolved question in this context is whether parts of the type IV pilus machinery such as the minor pilins and the pilin PilA act as piston to push the slime out of the PilQ gate as has been suggested (88). The identification of PilQ/GspD as slime nozzle is therefore a necessary first step to allow testing these various hypotheses on the contributions of slime secretion to motility in these various bacteria. In this context, our observations of GspD-depleted cells clearly demonstrate that slime secretion contributes to gliding motility in *M. xanthus*. Thus, we provide direct molecular evidence that slime contributes to motility, and identify *gspD* as a *bona fide* A-motility gene. Moreover, that *gspD* is essential also explains why the nozzle has so far never been identified in genome-wide genetic screens (77), and suggests the possibility that additional key components of A-motility remain to be found.

Alone, our results do not address the debate about the role of slime secretion in A-motility, since all current models propose a requirement for slime secretion. If slime secretion provides the propulsive force for motility, cells lacking slime secretion should lack A-motility, but the same would be true if slime is an important adhesion that provides surface contacts necessary for other molecular motors to act on (76). Therefore, additional experiments are required to address the precise role of slime secretion in A-motility; for example, the analysis of the chemical composition of the slime and its physicochemical properties, the identification and deletion of genes involved in its synthesis, and the determination of whether cells must themselves secrete slime to be motile, or simply require slime in their environment. Equally important will be to answer how widespread is the use of secretins as high-through-put nozzles for EPS secretion in Gram-negative bacteria.

## Materials and methods

### Bacterial Strains and Growth Conditions

*M. xanthus* cells were grown in CTT (1% casitone, 10 mM Tris pH 8.0, 8 mM MgSO_4_, 1 mM KH_2_PO_4_) or ½ × CTT (0.5% casitone, 10 mM Tris pH 8.0, 8 mM MgSO_4_, 1 mM KH_2_PO_4_) and maintained on CTT plates with 1.5% agar (65). When appropriate, 100 μg/ml kanamycin or 15 μg/ml oxytetracycline was used for selection. *A. platensis* strain LB 2340 from the Texas Algal Culture collection UTEX was grown under constant white light using an alkaline *Spirulina* medium: solution I (162 mM NaHCO_3_, 38 mM Na_2_CO_3_, and 2.9 mM K_2_HPO_4_ in 500 ml dH_2_O) and II (29.4 mM NaNO_3_, 5.74 mM K_2_SO_4_, 17.1 mM NaCl, 0.81 mM MgSO_4_, 0.27 mM CaCl_2_ in 500 ml dH_2_O) were autoclaved separately, combined after cooling, and 2 ml of a sterile-filtered 0.1 mM vitamin B_12_ solution was added. The freshwater cyanobacteria *O. lutea* (SAG 1459-3) and *Ph. autumnale* (strain Chesterfield; isolated by Dr Aya Farag from the University of Sheffield from a drainage site in Chesterfield and identified by 16S rRNA sequencing) were grown in BG11 medium (17.6 mM NaNO_3_, 0.23 mM K_2_HPO_4_, 0.3 mM MgSO_4_, 0.24 mM CaCl_2_, 0.031 mM citric acid, 0.021 mM ferric ammonium citrate, 0.0027 mM Na_2_EDTA, 0.19 mM Na_2_CO_3_, 1 ml trace metal mix in 1000 ml dH_2_O). Both strains, *O. lutea* and *Ph. autumnale* were sequenced by MicrobesNG (Birmingham). Strains used are listed in **Table 1**.

**Table 1.** Bacterial strains used in this study

| **<u>Strain</u>** | **<u>Relevant description</u>** | **<u>Source or reference</u>** |
| --- | --- | --- |
| *Arthrospira platensis* |  |  |
| LB 2340 | Wild type | (108) |
| *Myxococcus xanthus* |  |  |
| DK1622 | Wild type | (65) |
| DK10407 | [DK1622] $\Delta pilA$ | (109) |
| DK8615 | [DK1622] $\Delta pilQ$ | (66) |
| EH098 | [DK1622] $\Delta gspD$ P<sub>cuoA</sub>:*gspD*. Deletion of *gspD* at the chromosomal locus with complementation at the chromosomal *attB* site under control of the copper-inducible *cuoA* promoter. Tet<sup>R</sup>. | This study |
| EH099 | [DK10407] $\Delta pilA$ $\Delta gspD$ P<sub>cuoA</sub>:*gspD*. Deletion of *gspD* at the chromosomal locus with complementation at the chromosomal *attB* site under control of the copper-inducible *cuoA* promoter in a social-motility defective background. Tet<sup>R</sup>. | This study |
| *E. coli* |  |  |
| BL 21 Star (DE3) | *E. coli* host for protein expression. | Invitrogen |
| TOP10 | *E. coli* host for plasmid maintenance. | Invitrogen |

### Construction of Copper-inducible Mutants

To generate a markerless deletion of the *gspD* gene, we transformed the parent cell line with pDMZ96 (**Table 2**) and selected for plasmid integration with 100 μg/ml of kanamycin. Clones were selected and grown in media lacking kanamycin, plated in media containing 2.5% galactose to select for loss of the plasmid, and screened by PCR for gene deletion. Multiple attempts to delete *gspD* in several genetic backgrounds failed; consistent with the conclusion that *gspD* is an essential gene. As a secondary strategy, clones that had integrated the deletion plasmid were transformed with plasmid pDMZ94, which expresses the *gspD* gene regulated by the copper inducible promoter *P_cuoA_* from the Mx8 phage attachment site (69). Clones were collected and grown in media containing 300 μM CuSO_4_ and subject to galactose selection. Multiple clones containing the *gspD* deletion at the native chromosomal locus were recovered, and maintained in CTT media supplemented with 300 μM CuSO_4_.

**Table 2.**
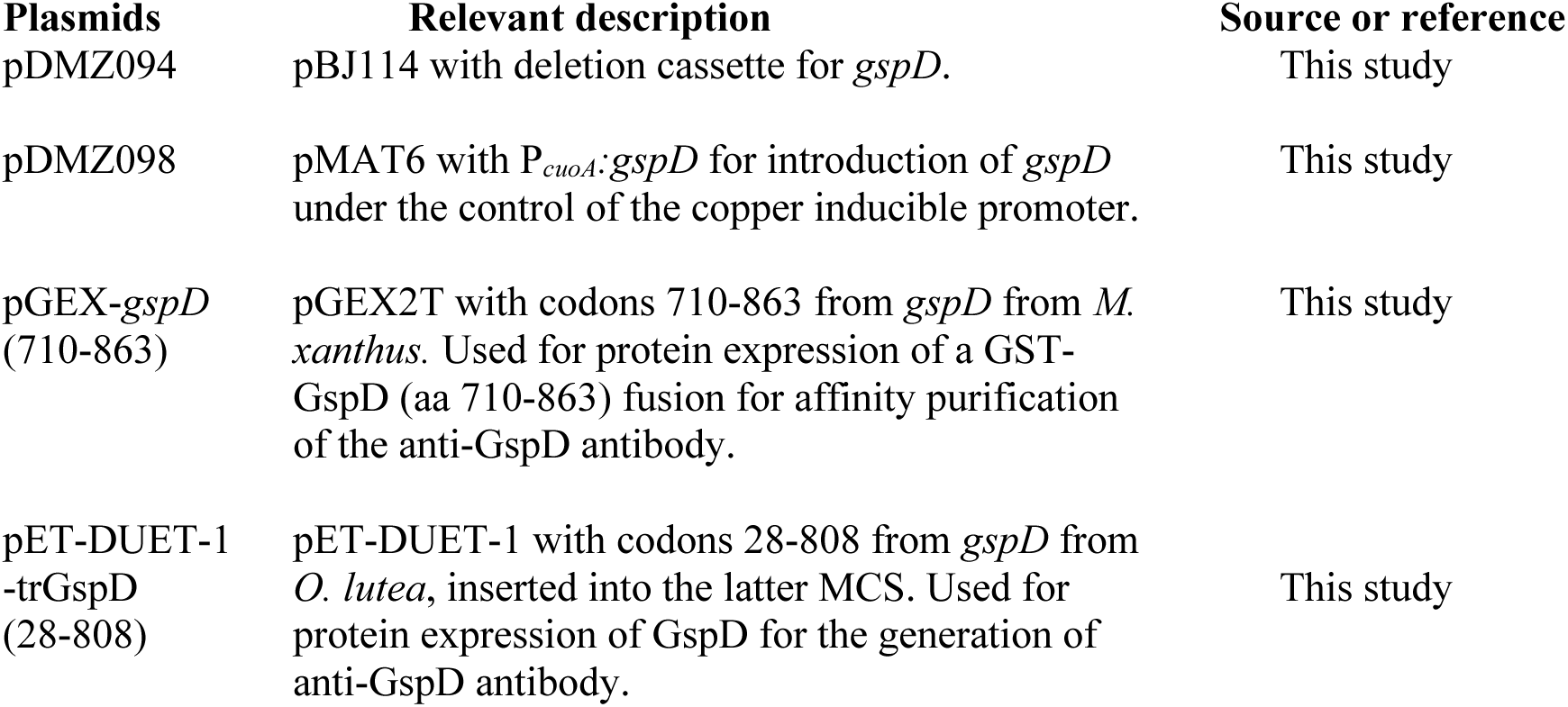
Plasmids used in this study

### Isolation and Purification of PilQ/GspD Nozzles

To isolate nozzles from the three cyanobacteria species, ca. 100 g wet weight of cells were harvested by centrifugation (10 min at 1,000 × g), washed twice in Tris-HCl buffer (10 mM Tris-HCl, pH 7.5), and chilled on ice. Cells were disrupted by glass beads using a Desintegrator S cell mill (Bernd Euler Prozesstechnik, Frankfurt) at 0 °C and unbroken cells were removed by low speed centrifugation (10 min at 1,000 × g). Crude cell envelopes were collected on ice and further purified using a Percoll density gradient (15% vol/vol) for 1 h at 10,000 × g. The pale orange-colored pellet at the bottom of the gradient contained highly enriched cell envelopes. After several washes with Tris buffer, the purified envelopes were re-suspended in the buffer containing 2% Triton X-100 and 0.02% sodium azide. The suspension was shaken overnight at 37 °C and the autolytic digestion of the peptidoglycan monitored by light microscopy. Undigested cross walls and debris were removed by centrifugation (10 min at 50,000 × g) and crude nozzle preparations were collected in the ultracentrifuge (1 h at 366,000 × g) before being further purified using a CsCl density gradient (0.3 g/ml). After overnight centrifugation, the band containing the nozzles was collected using a gradient fractionator (Labconco Auto Densi-Flow), dialyzed against Tris buffer, and the nozzles were either collected by centrifugation (1 h at 366,000 × g) or further purified using 30 ml gradients of 10-40% sucrose (wt/wt). One milliliter of the nozzle-containing suspension was dialyzed against Tris buffer and then layered on top of the gradient and centrifuged at 100,000 × g for 17 h using a Beckman SW41 rotor. The twelve collected fractions were dialyzed against Tris buffer, examined in the electron microscope for the presence of nozzles using carbon-coated copper grids that were either glow discharged or not, and analyzed using SDS-PAGE. Proteins were identified using Edman degradation and mass spectrometry.

To isolate GspD nozzles from *M. xanthus*, ca. 80 g wt or Δ*pilQ* cells were collected by centrifugation and re-suspended in 1 M sucrose by vigorous shaking. Cells and cell debris were removed by differential centrifugation (17,000 × g for 10 min followed by 32,000 × g for 10 min), and five volumes of chilled Tris buffer were added to dilute the sucrose. Enriched OMs were pelleted by centrifugation (10 min 50,000 × g) and re-suspended in Tris buffer at a concentration of 0.1 g/ml. An equal volume of 1% solution of dodecyl-maltoside was added to solubilize the OMs and un-solubilized material removed by centrifugation (10 min 50,000 × g). After addition of 0.3 mg/ml CsCl, the solution was centrifuged overnight at 366,000 × g using a Beckman SW 55 Ti rotor. A turbid yellowish band was visible about 2/3 of the way in the gradient and was identified as enriched in nozzle-like structures by TEM. These nozzle-containing bands were harvested, dialyzed against Tris buffer, and either directly analyzed or further purified as described above for the cyanobacteria.

### Antibody Production

His-GspD and His-PilQ_Olut_ were expressed in *Escherichia coli* BL21 cells and purified according to the manufacturer’s instructions (Invitrogen, Carlsbad, CA). The proteins were injected into rabbits to generate polyclonal antibodies according to standard protocols (His-GspD, Cocalico, Reamstown, PA; His-PilQ_Olut_, Eurogentec, Seraing, BE). Sera were tested for cross-reactivity by immunoblotting lysates from wildtype *M. xanthus* or cyanobacterial cells. To increase the specificity of the reactivity, we affinity purified the His-GspD antibodies. Amino acids 710-863 of GspD were expressed as a C-terminal fusion to the glutathione S-transferase protein (GST-GspD C-term) in *E. coli* BL21 strain, and captured with glutathione sepharose beads (GE Healthcare, Laurel, MD). Protein was eluted with 10 mM glutathione in 50 mM Tris, pH 8.0, 5% glycerol, and examined for purity by SDS-PAGE and Coomassie staining. Five hundred micrograms of protein were dialyzed against binding buffer (PBS with 10 mM EDTA) and re-bound to glutathione sepharose beads. Protein was then crosslinked to beads with 5 mg/ml DTSSP (Thermo Fisher Scientific, Rockville, MD) in binding buffer for 45 min at RT. Buffer was drained and the reaction quenched by washing beads twice for 5 min with 100 mM Tris, pH 8.0. The beads were then washed extensively with binding buffer, and elution buffer (4 M MgCl_2_) to remove any un-crosslinked protein. Beads were normalized with binding buffer, and incubated with the antisera overnight at 4 °C. Sera were drained, and beads washed twice with wash buffer (10 mM Tris, pH 7.5, 0.2% deoxycholic acid) and twice with wash buffer plus 0.5 M NaCl. Bound antibody was eluted with elution buffer, and collected in 1 ml fractions in tubes containing 50 μl of 10 mg/ml bovine serum albumin, and transferred immediately to dialysis bags and dialyzed against 1 L of PBS plus 0.02% sodium azide. Antibodies were tested for activity by immunoblot against lysates from *M. xanthus* or nozzle-enriched fractions from the cyanobacterial cell envelope preparations and recognized a single band. Affinity purification was not necessary for the cyanobacterial antibody as it recognized only a single band in our species.

### SDS-PAGE and Immunoblotting

Equal cell numbers from liquid grown cultures or equal amounts of CsCl fractions were solubilized in 2× Tris-Glycine SDS buffer (Life Technologies) by boiling for 15 min. Samples were separated by SDS-PAGE and transferred to a PDF membrane (Millipore, Billerica, MA). The membrane was blocked with PBS containing 0.5% tween (PBST) and 5% milk, and probed overnight with affinity purified anti-GspD or anti-PilQ in PBST plus 3% BSA. The membrane was washed with PBST, and probed with HRP-conjugated anti-rabbit antibody (Jackson ImmunoResearch, West Grove, PA) in PBST containing 5% milk. HRP was activated using SuperSignal West Pico Chemiluminescent Substrate (Thermo Scientific, Rockford, IL) and imaged with a FluorChem Q system (Protein Simple, Wallingford, CT).

### Electron Microscopy

To visualize slime secretion, carbon-coated gold grids (EMS) were glow discharged, coated with acid-hydrolyzed chitosan (54), and dried. Grids were held face-up by forceps, and 2 μl of a suspension of cells grown in the absence or presence of copper were spotted onto the grid. Cells were incubated at room temperature (RT) in a humidity chamber for 20 min. Grids were rinsed with H_2_O and routinely stained with 1.5% silico phosphotungstate (SiPTA), pH 7.4 or, to identify non-slime material, with un-buffered UA (pH 4.5) or SiPTA, pH 4.0 citric acid. Grids were examined with a Hitachi 7600 or a Philips CM120 transmission electron microscope at 80 kV, and micrographs collected using AMT Image Capture Engine software controlling an AMT ER50 5 megapixel CCD camera (Advanced Microscopy Techniques Corp., Danvers, MA).

To quantify the number of slime trails per cell, EM grids were prepared as above using cells grown for 24 h in liquid media containing 0.01, 0.2, or 0.5 mM CuSO_4_. Prepared grids were examined by EM, and isolated cells (>1 full cell-length from nearest neighboring cell) were selected at low magnification, so that slime trails could not be observed prior to imaging (to reduce experimenter bias). High magnification images were collected, and the numbers of slime trails emanating from at least 12 cells/condition were counted. Cells with disrupted OM were excluded. The average length of cells did not significantly vary between the populations (determined by one-way ANOVA (mean ± S.D.: 0.01 mM CuSO_4_ = 9.4 ± 2.4 μm; 0.2 mM CuSO_4_ = 11.5 ± 4.2 μm; 0.5 mM CuSO_4_ = 9.2 ± 2.4 μm)). Data are presented as the mean number of slime trails per cell, with the standard deviation.

To disrupt OM for visualization of nozzles, cells swarming on hard agar with or without copper were scraped into CTT media in a 1.5 ml centrifuge tube. An equal volume of 710-1180 μm glass beads was added (Sigma-Aldrich, St. Louis, MO), and samples were subjected to vortexing at maximum power for 2 min. Cells were applied to a glow discharged EM copper grids, stained with 2.5% UA and imaged as above.

Cryosubstitution of cyanobacterial cells was performed as described (55). Briefly, *A. platensis* cells were high-pressure frozen using a Leica EM PACT2 instrument (Leica Microsystems, Buffalo Grove, IL), cryo-substituted for 80 h at -87 °C in acetone containing 2% osmium tetroxide, and, after slowly warming to RT, embedded in Spurr’s resin. Thin sections were stained with UA and lead nitrate (104), and examined in a Philips CM12 electron microscope. To visualize nozzles in membranes, isolated outer membranes were picked up on 200 mesh carbon-coated copper grids and unilaterally shadowed with Pt/C at an angle of 45°. Images at various magnifications were recorded as described above.

### Immunoelectron Microscopy of Isolated GspD Nozzles

CsCl gradient fractions containing cell envelope proteins of *A. platensis* were adsorbed for 15 sec to 200 mesh carbon-coated gold grids, washed with water and PBS and then incubated for 40 min with a 1:500 dilution of a serum from a rabbit inoculated with GspD from *M. xanthus*. The grids were washed on three drops of PBS before being incubated for 12 min with a 5 nm gold-labelled anti-rabbit secondary antibody (Jackson ImmunoResearch West Grove, PA) at dilutions of 1:10. After repeated washes with PBS and water, the grids were stained with 2% un-buffered UA and viewed under the electron microscope. As negative control, anti-BacM rabbit serum was used. To judge labelling, 200 randomly selected PilQ complexes of the sample and the negative control were scored for the presence of gold label.

### Image Analysis and Particle Averaging

A total of 1605 single, double, or multiple PilQ complexes were selected using PyTom, classified through iterative multivariate statistical analysis (MSA), and aligned using a single reference dimer particle (105). For MSA, twelve eigenvectors were used to classify the particles into four separate classes, which were then aligned and averaged using the TOM toolbox programs (106).

### Immunofluorescence (IF) Light Microscopy

*M. xanthus* were grown 24 h in the absence or presence of 300 μM CuSO_4_ and adhered to sterile glass coverslips overnight in CTT media, with or without copper. Cells were then processed essentially as described (107). Briefly, cells were rinsed with PM buffer (20 mM Na-phosphate, 1 mM MgSO_4_, pH 7.4) and fixed with 4% paraformaldehyde in PM buffer. Cells were permeabilized with 0.2% Triton X-100 and 1 mg/ml lysozyme, and probed with affinity purified anti-GspD antibody at a 1:10 dilution in PBS buffer with 2% BSA. Secondary antibody was Alexa_594_-conjugated anti-rabbit (Life Technologies, Carlsbad, CA) diluted 1:1000 in PBS with 2% BSA. Cells were stained with 1 μg/ml DAPI, and examined with a Nikon Eclipse 90i microscope with a 100×/NA 1.4 phase-contrast oil immersion objective (Nikon, Melville, NY). Images were collected with an ORCA ER CCD camera (Hamamatsu, Bridgewater, NJ) and processed using Volocity (PerkinElmer, Waltham, MA).

For *O. lutea*, actively growing and motile cell filaments were collected, washed in ddH_2_O, and left at 4°C overnight to allow autolysis. For *Ph. autumnale*, actively growing and motile cell filaments were collected, washed in BG11 medium, and incubated at 50 °C for 14 hours. For both, cells were then treated with 0.2 M Glycine buffer at pH 2.5 for 15 min at RT. After thorough washing in 20 mM HEPES at pH 8, cells were air-dried onto poly-L-lysine (PLL)-coated coverslips and submerged in 70% ethanol at -25 °C for 30 minutes for fixation. Coverslips were washed in PBS thoroughly and blocked in PBS containing 2% BSA and 0.5% Tween-20 at RT. Coverslips were placed cell-face down onto 100 μl drops of primary anti-PilQ at 1:600 dilutions in blocking buffer at RT. After one hour, coverslips were washed in blocking buffer, and further labelled with Alexa Fluor 488-conjugated secondary antibodies (Invitrogen, Carlsbad, CA) for one hour at RT as above. After washing, coverslips were mounted with SlowFade Gold antifade mountant (Molecular Probes, Carlsbad, CA) and sealed with nail polish. Imaging was performed with a Nikon Eclipse Ti inverted fluorescence microscope using the Nikon Plan Apo 100× Ph oil (NA 1.45) objective. This was equipped with the Andor Zyla sCMOS camera (Andor, Belfast, NI). Image acquisition was controlled using NIS Elements AR 4.2 imaging software (Nikon Instruments, Netherlands). Images were visualized and analyzed with FIJI (110).

### Fluorescence Imaging of EPS Secretion

Actively motile *O. lutea* or *Ph. autumnale* cell filaments were collected and washed in BG11 media. The filaments were subjected to brief sonication (<1 second) at low power or cut into short fragments using razor blades. They were transferred to an ice-cold solution of BG11 with 10µg/ml of Alexa Fluor 488-congujated concanavalin A (Invitrogen, Carlsbad, CA). Imaging was performed in a temperature regulated chamber set to 28°C, using the same microscope for the IF imaging of *O. lutea* and *Ph. autumnale* described above. Cells were seeded into an ibidi PLL-treated µ-Slide VI^0.4^ flow channel slide (ibidi, Gräfelfing, Germany) and allowed to settle for 5 minutes. BG11 was flowed through at >0.5 ml/s to remove excess concanavalin A, and to encourage dissociation of slime bands from cell surfaces.

### Serial Dilution Growth Assay

To test the requirement for copper for growth of the *M. xanthus* strains, cells were grown overnight in CTT media with 200 μM CuSO_4_ at 32 °C. Cells were sub-cultured into CTT media with lacking copper, but with 200 μM of the copper chelator bathocuproinedisulfonic acid (BCS, Sigma-Aldrich, St. Louis, MO) and grown for 24 h at 32 °C. These cells were again sub-cultured into media with 200 μM BCS and grown for an additional 24 h at 32 °C. Cells were concentrated to 1 × 10^9^ cells/ml in CTT and four 4-fold serial dilutions were prepared. Three microliters of each cell suspension were spotted on CTT plates with 1.5% agar, containing either 200 μM BCS, 100 μM CuSO_4_, or 500 μM CuSO_4_, dried, and incubated at 32 °C for 48 h.

### Motility Assays

For adventurous motility assays, mutant *M. xanthus* cells were grown overnight in liquid culture containing 200 μM CuSO_4_ at 32 °C. Cells were then sub-cultured into media containing 200 μM BCS and grown for 24 h at 32 °C. Wildtype and Δ*pilA* cells were grown overnight in the absence of copper. Cells were diluted to 1 × 10^8^ cells/ml in CTT, and 10 μl were spotted onto 1/2× CTT plates with 1.5% agar, and either 200 μM BCS or 300 μM CuSO_4_. Spots were dried and plates were incubated for 48 h at 32 °C. Swarm edges were examined with a Nikon Inverted TE200 microscope, using a 10× objective, and digital images were collected with a SPOT RT camera and SPOT Basic software (Diagnostic Instruments, Inc., Sterling Heights, MI).

## Acknowledgments

We thank the W. Harry Feinstone Department of Molecular Microbiology and Immunology at the Johns Hopkins Bloomberg School of Public Health for generous support during the initial phase of this investigation, members of the Hoiczyk laboratory for helpful discussions and comments on the work and the manuscript, Colleen McHugh and Harriet Pratley for the generation of GspD/PilQ antisera, José Muñoz-Dorado for providing plasmids for copper-inducible gene expression, Daniel Bollschweiler, Florian Beck, and Harald Engelhardt (MPI of Biochemistry, Martinsried) for their help with averaging the isolated *A. platensis* GspD complexes, Christopher Hill for help with the immunogold labeling and high pressure freezing, and Carolyn Machamer for providing research space for D.M.Z. Mass spectrometry analyses were performed by the biOMICS/chemMS Facility of the Faculty of Science Mass Spectrometry Centre at the University of Sheffield (*O. lutea*, *Ph. autumnale* and *A. spirulina*) and the Mass Spectrometry & Proteomics Resource Core at Harvard University (*A. spirulina*). This research was funded in part by a National Institutes of Health General Medicine Grant (GM85024 to E.H.), a Forschungsstipendium of the Max Planck Society (to E.H.), a National Institutes of Health Infectious Disease and Immunology Training Grant (to D.M.Z.), and a National Science Foundation Grant (1949762 to D.M.Z.). In addition, E.H., J.M.T.S., and D.M.Z. acknowledge support from the Imagine: Imaging Life initiative of the University of Sheffield.

---

**Figure S1.**
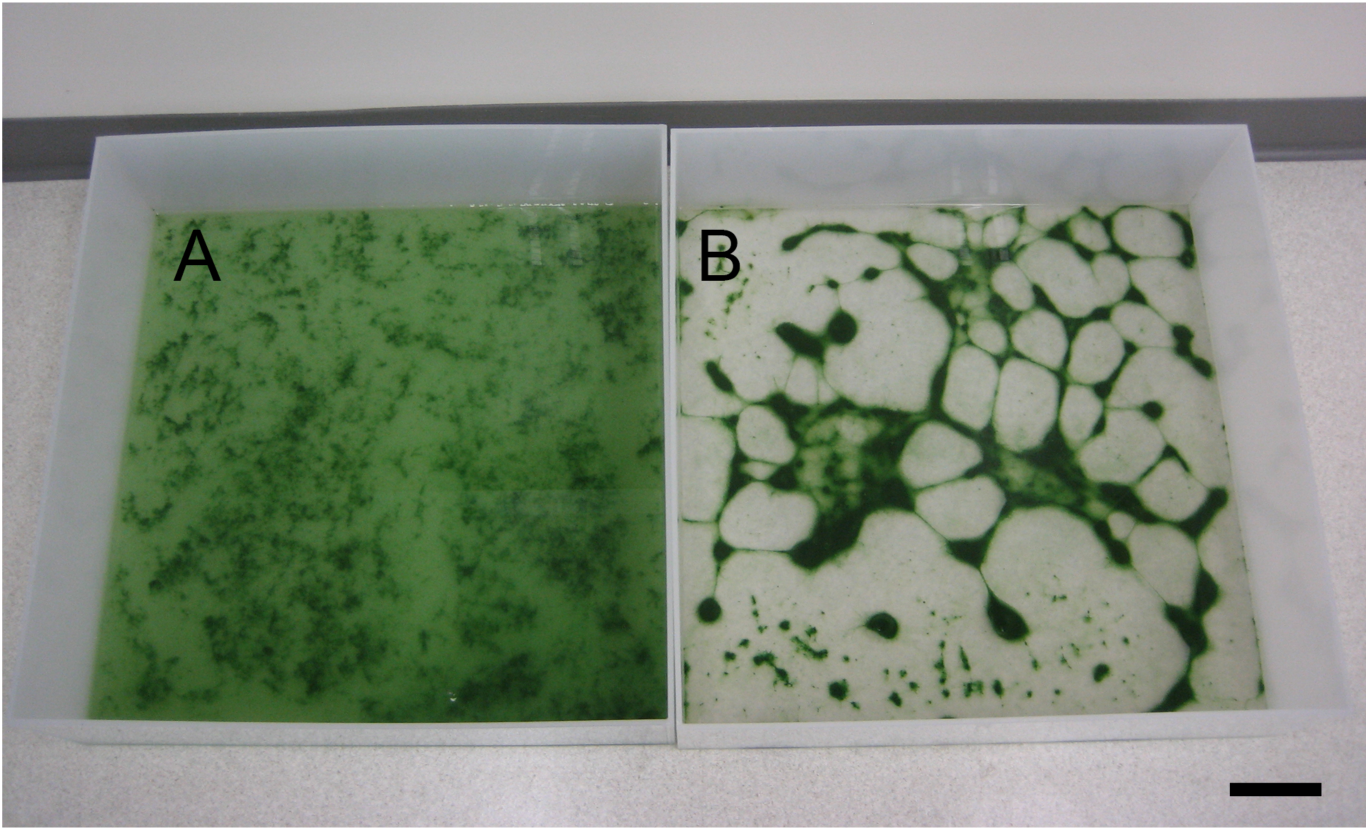
Addition of cAMP to a culture of *Arthrospira platensis* (A) induces motility-dependent clumping (B), revealing the ability of the cells to glide. As motility-dependent clumping is invariably correlated with slime secretion and nozzle presence in this species, clumping can be used to screen for the presence of nozzles. Scale bar, 10 cm.

**Figure S2.**
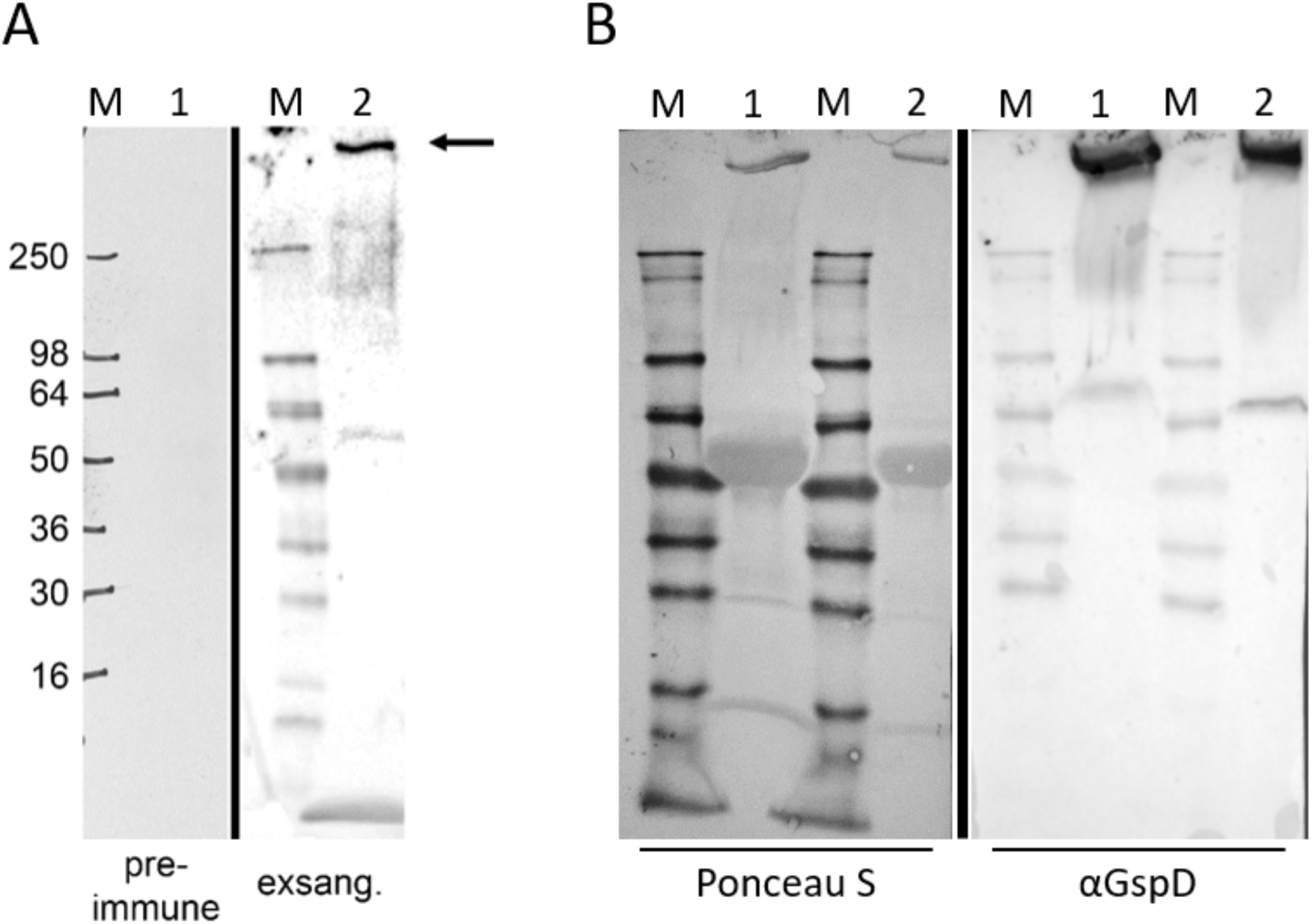
Antibodies raised against GspD from *Myxococcus xanthus* and PilQ from *Oscillatoria lutea* specifically recognize PilQ from *Arthrospira platensis,* and *Oscillatoria lutea* and *Phormidium autumnale,* respectively. (A) *A. platensis* cell envelopes were isolated, solubilized, and fractioned using CsCl gradients. Protein samples were separated by SDS-PAGE, transferred to a membrane and probed with serum from a rabbit inoculated with GspD from *M. xanthus*. The >250 kDa band of PilQ from *A. platensis* (see Fig. 1G) was the only band to react with the antibody. As a negative control, *A. platensis* total cell protein was separated and probed with an identical dilution of pre-immune serum resulting in no detectable signal. Note, the left panel is recorded using HyBlot CL autoradiography film (Denville, Metuchen, NJ), while the right panel is generated with the ChemiDoc Imaging System (BioRad, Hercules, CA). (B) Roughly equal amounts of PilQ-nozzle-enriched CsCl gradient fractions of *O. lutea* and *Ph. autumnale* nozzle preparations were dialyzed, boiled in sample buffer, and separated on two SDS-PAGE gels. One of the gels was stained with Coomassie G250 and the >250 kDa large band of PilQ analyzed by mass spectrometry (see Table S1 for mass spec data). The second gel was blotted onto nitrocellulose and imaged with a FluorChem Q system (Protein Simple, Wallingford, CT) after staining with Ponceau S (left panel). Following de-staining, the membrane was probed with serum from a rabbit inoculated with PilQ from *O. lutea* (expressed in and purified from *E. coli*) and imaged with the FluorChem Q system. Lane 1, *O. lutea*; lane 2 *Ph. autumnale*. Similar to Fig. S2A, the pre-immune serum did not result in a detectable signal (not shown).

**Figure S3.**
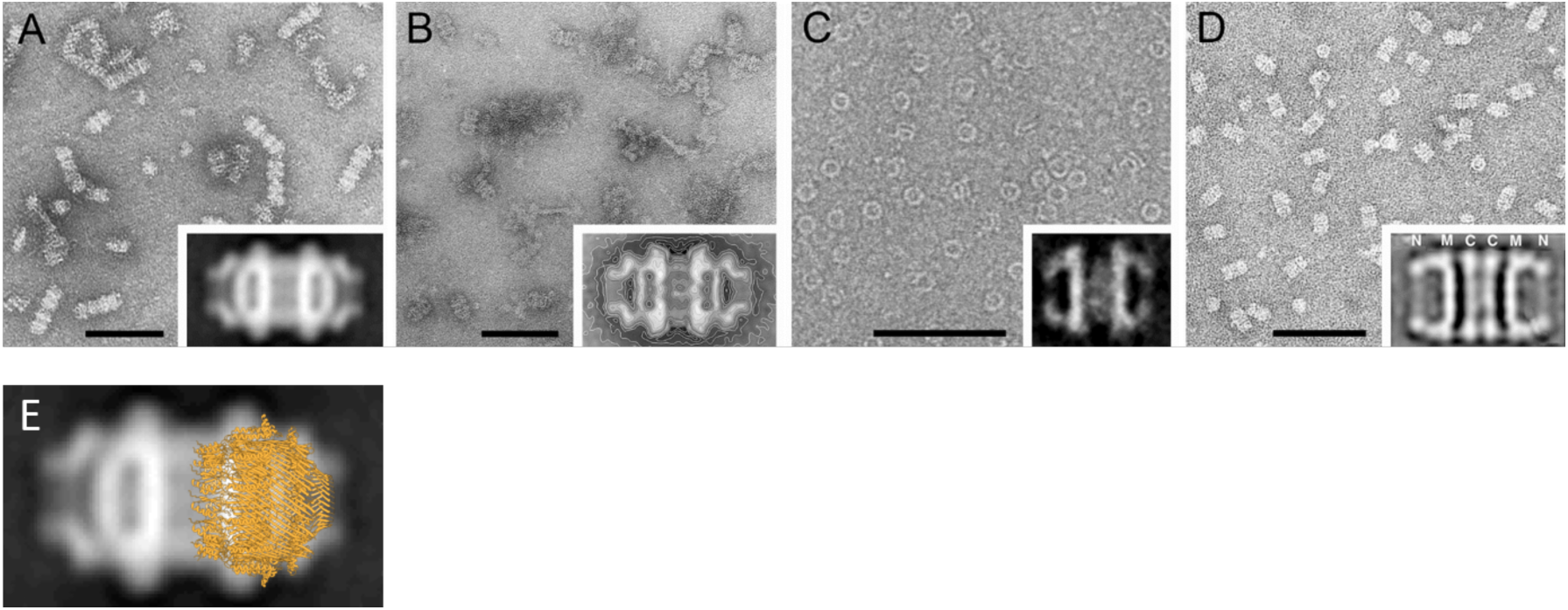
Comparison of selected isolated PilQ dimer complexes following negative staining and averaging. Despite large variations in the molecular masses of PilQ monomers and of the overall length of the complexes (*Arthrospira platensis* PilQ: 756 aa, 81 kDa, 32nm length; *Phormidium uncinatum* PilQ: molecular weight unknown, 32 nm length; *Klebsiella oxytoca* PulD^28-42/259-660^: 416 aa, 44 kDa, length 15 nm; filamentous phage f1, secretin pIV: 405 aa, 43 kDa, length 24 nm; all amino acid data and mass calculations correspond to the mature proteins lacking signal peptides), all isolated dimers show similar, characteristic structural features, further supporting the idea that the cyanobacterial complexes are formed by PilQ. These structural features comprise large openings that provide access to a vestibule-like chamber whose bottom is formed by a massive ring structure, followed either by a narrow ring (pIV complex) or a poorly structured constriction (PilQ dimers of cyanobacteria and *K. oxytoca*). (A) Isolated and enriched PilQ dimer complexes from the cyanobacterium *A. platensis*. The multimeric complexes are often arranged in chains of two, three, or, rarely, greater numbers of complexes. The inset shows the characteristic appearance of the 32 nm-long dimers, based on the two-fold symmetrized average of 902 particle halves. (B) Isolated PilQ dimers of the filamentous cyanobacterium *Ph. uncinatum* (51). The averaged dimers have the same overall dimensions (32 × 20 nm) and architecture as the PilQ dimers of *A. platensis*, revealing that these structures are identical. The inset shows the average of 334 side views. (C) Isolated native PulD complexes from the bacterium *K. oxytoca* (1). In contrast to the cyanobacterial PilQ complexes, only monomeric rings ca. 13 nm wide and 10 nm long were isolated. Replacement of a highly-conserved threonine with isoleucine (T470I) or valine (T470V) in a partially shortened version of PulD^28-42/259-660^ resulted in the formation of a small subpopulation of ca. 15 nm long dimers (2). The inset shows the average of 16 such T470I dimer complexes. (D) Isolated pIV secretin dimer complexes from the filamentous phage f1 (12). Similar to the PilQ complexes of *A. platensis* and *P. uncinatum*, pIV complexes were recovered as dimers during the isolation, a result that was suggested to be due to the use of detergent during the purification process. The letters N, C, and M denote the suggested positions of the N- and C-termini and the middle part of the monomers within each half particle, respectively. Note, that insets are not drawn to scale, but presented to demonstrate the morphological similarities between these complexes. All scale bars, 100 nm. (E) Overlay of the atomic structure of the *Vibrio cholera* GspD monomeric ring (PDB accession code 5WQ8) onto the 32 nm-long two-fold symmetrized dimers of the *A. platensis* PilQ structure. Note, the *A. spirulina* structure has a maximal width of 20 nm, while the *V. cholera* structure is only 16 nm wide. The image demonstrates that the isolated cyanobacterial PilQ structures are likely dimers of monomeric secretin rings as seen in *V. cholera* or *M. xanthus*.

**Figure S4.**
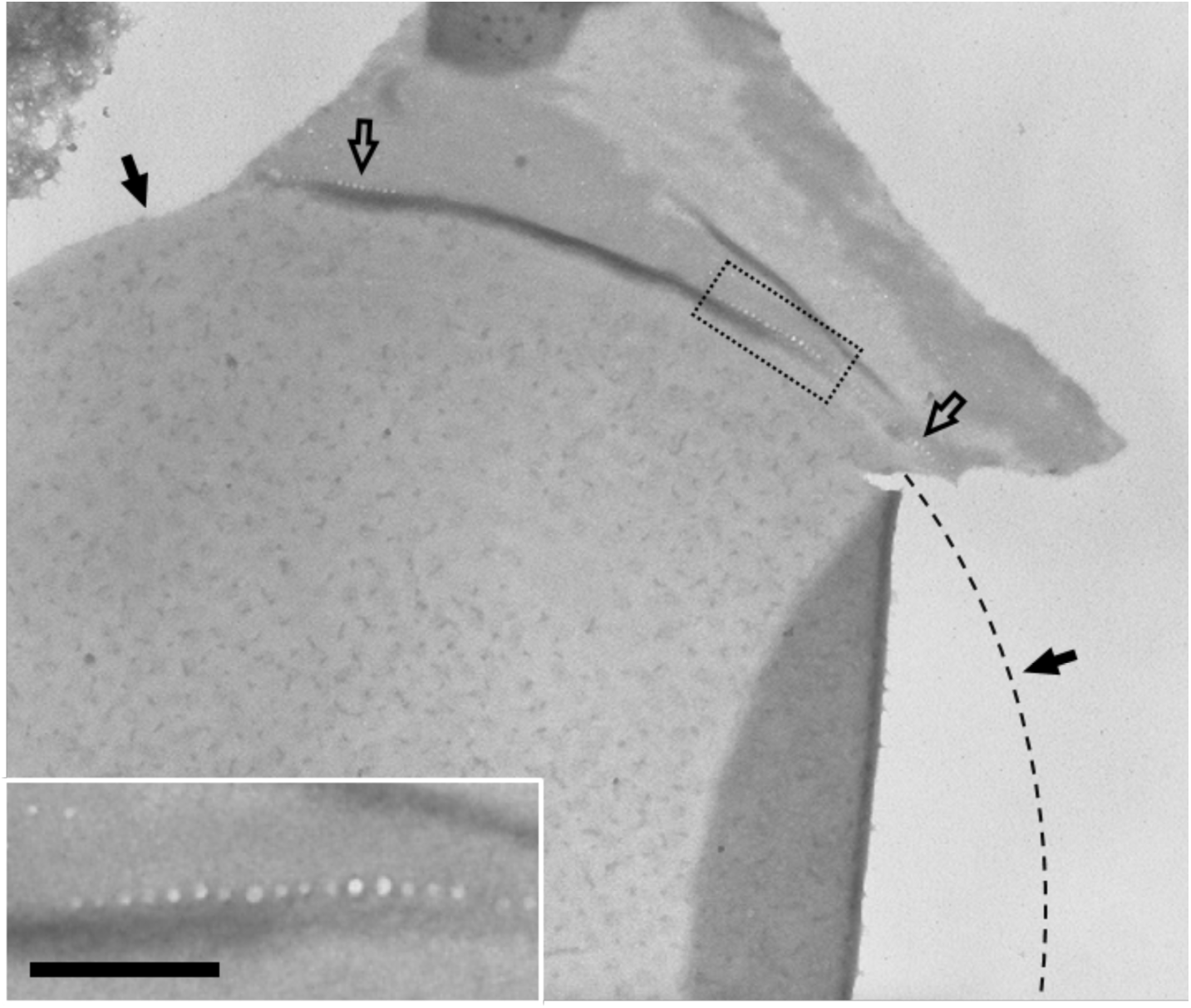
Disc-shaped cross wall of *Oscillatoria lutea* showing nozzle-containing portions of the attached longitudinal cell wall. Upon mechanical cell breakage and negative staining, individual disc-shaped cross walls can be found that still have either parts or the entire nozzle-containing longitudinal cell wall attached. The here visible incomplete presence of the nozzle-containing longitudinal cell wall explains why in the fluorescence light microscopy (Fig. 2A) the antibody often does not label the entire circumference of the disc-shaped cross wall. The solid arrows and the stippled line show the contour of the disc-shaped cross wall which at the right edge is partly folded over and therefore stained more darkly. The hollow arrows point to the parts of the attached longitudinal cell wall where the row of trans-peptidoglycan channels of the PilQ nozzle apparatus is visible. The inset shows a higher magnification of the boxed area of the nozzle-containing longitudinal wall clearly showing the row of channels on that side of the cross wall. Note, the other row of channels is on the opposite side of the cross wall and therefore only faintly visible if at all in the inset. Scale bar, 200 nm.

**Figure S5.**
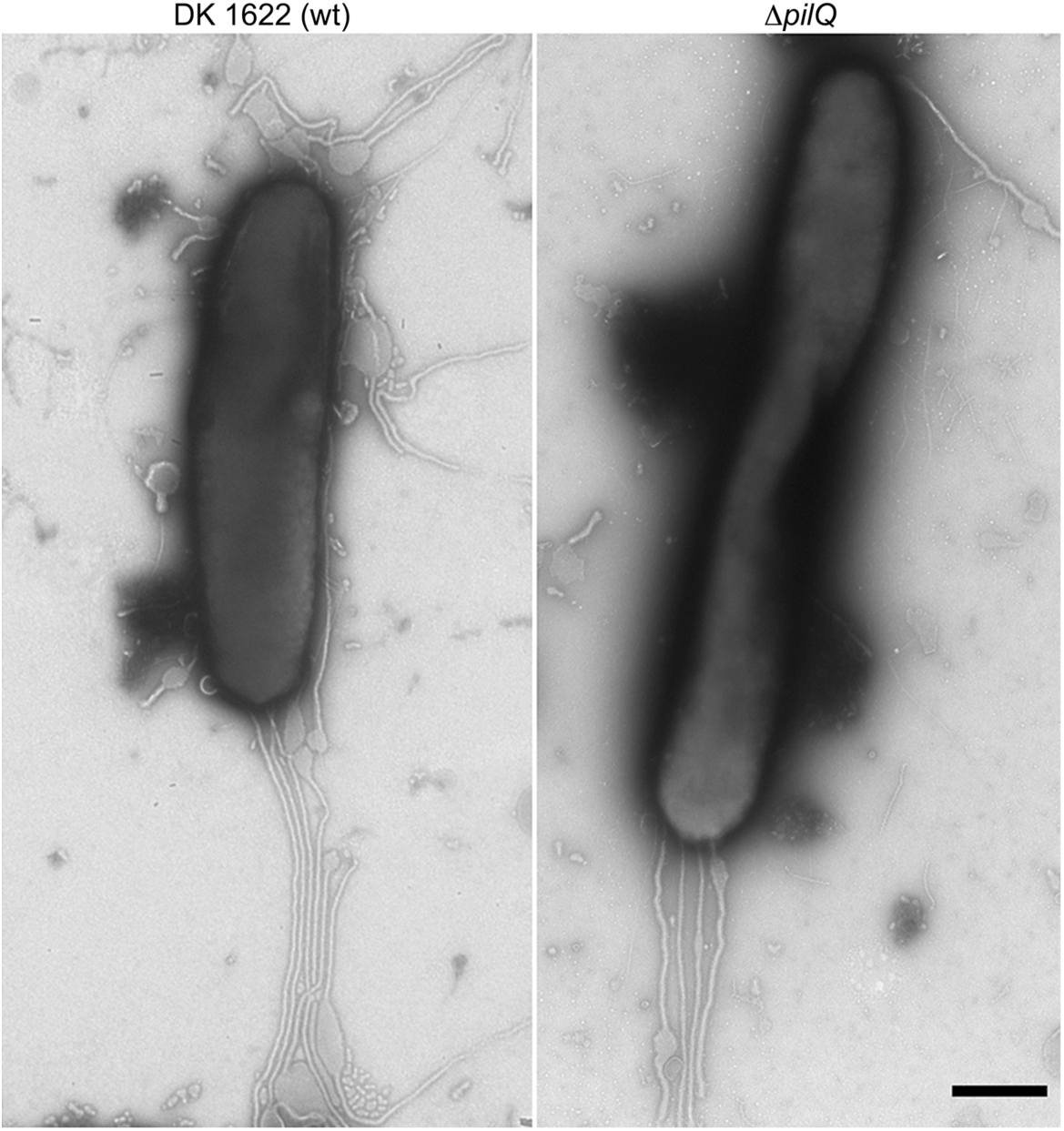
Slime secretion of *Myxococcus xanthus* Δ*pilQ* cells. To determine if the secretin PilQ is involved in slime secretion, wildtype and mutant strain cells were allowed to glide on chitosan-coated grids. Electron micrographs of representative cells show that the slime secretion of the two strains is indistinguishable confirming that only GspD engages in the secretion of slime. Scale bar, 500 nm.

**Figure S6.**
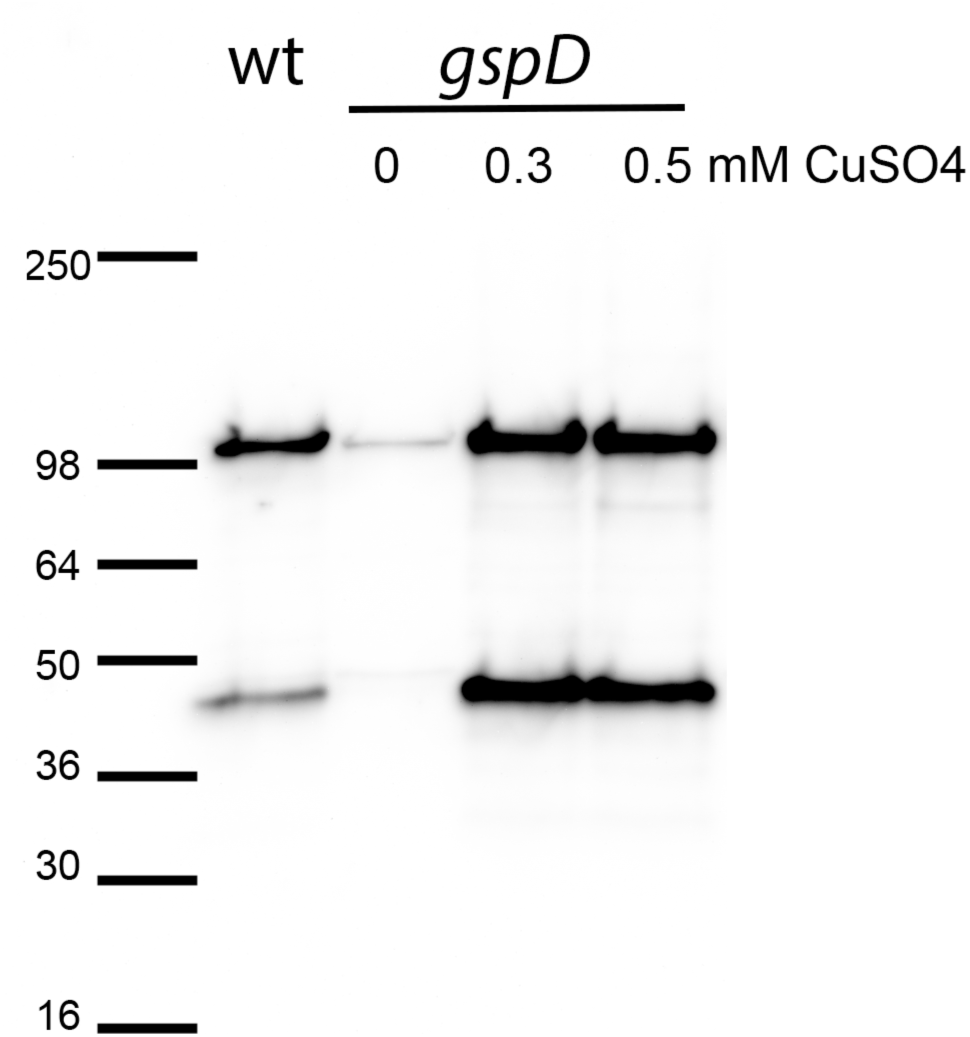
Copper-dependent overexpression of GspD in *Myxococcus xanthus*. Wildtype *M. xanthus* or mutants expressing *gspD* under control of a copper inducible promoter were grown with the indicated concentrations of CuSO_4_. Steady-state levels of GspD were evaluated by immunoblot with an affinity-purified anti-GspD antibody. This representative blot reveals that addition of copper to the medium results in an increase of *gspD* expression that eventually stabilizes at a copper concentration of about 0.3 mM. GspD is visualized as a ∼100 kDa band, and occasionally a second band at ∼47 kDa.

**Figure S7.**
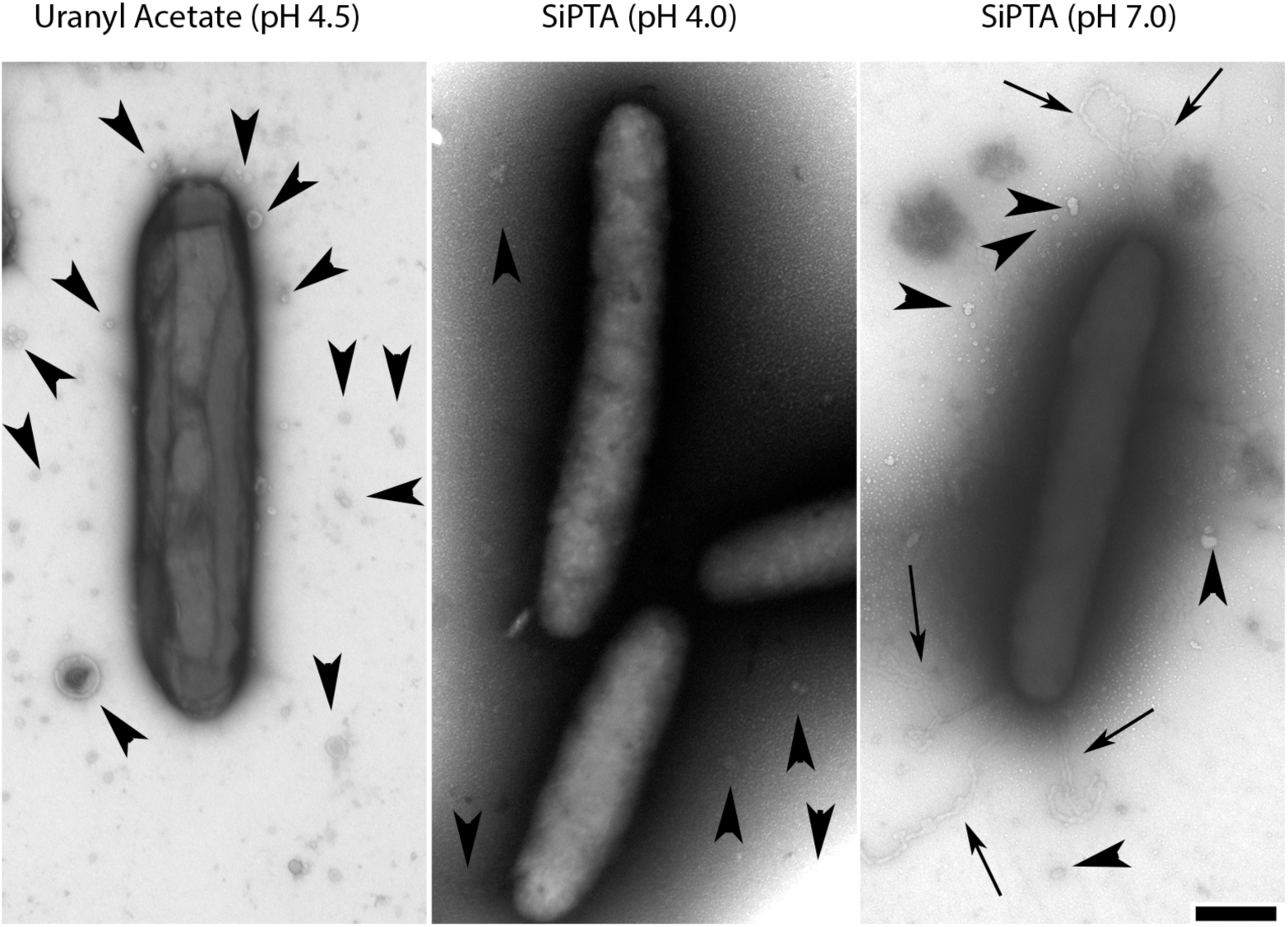
Identification and detection of slime trails of *Myxococcus xanthus* using electron microscopy of cells that are negatively stained at neutral pH. *M. xanthus* cells leave slime trails behind that have been visualized using numerous staining and microscopy methods. As these structures were originally discovered using TEM, we used their pH sensitivity during negative staining to discriminate them from other outer membrane-derived structures such as lipid tubules. Acidic stains such as un-buffered uranyl acetate (A) or silico phosphotungstic acid (SiPTA) (B) specifically remove the slime trails while SiPTA at neutral pH (C) preserves the slime trails. In contrast, membrane vesicles are not affected by the pH of the negative staining. Arrows, slime trails; arrowheads, extracellular vesicles; scale bar, 0.5 μm.

**Figure S8.**
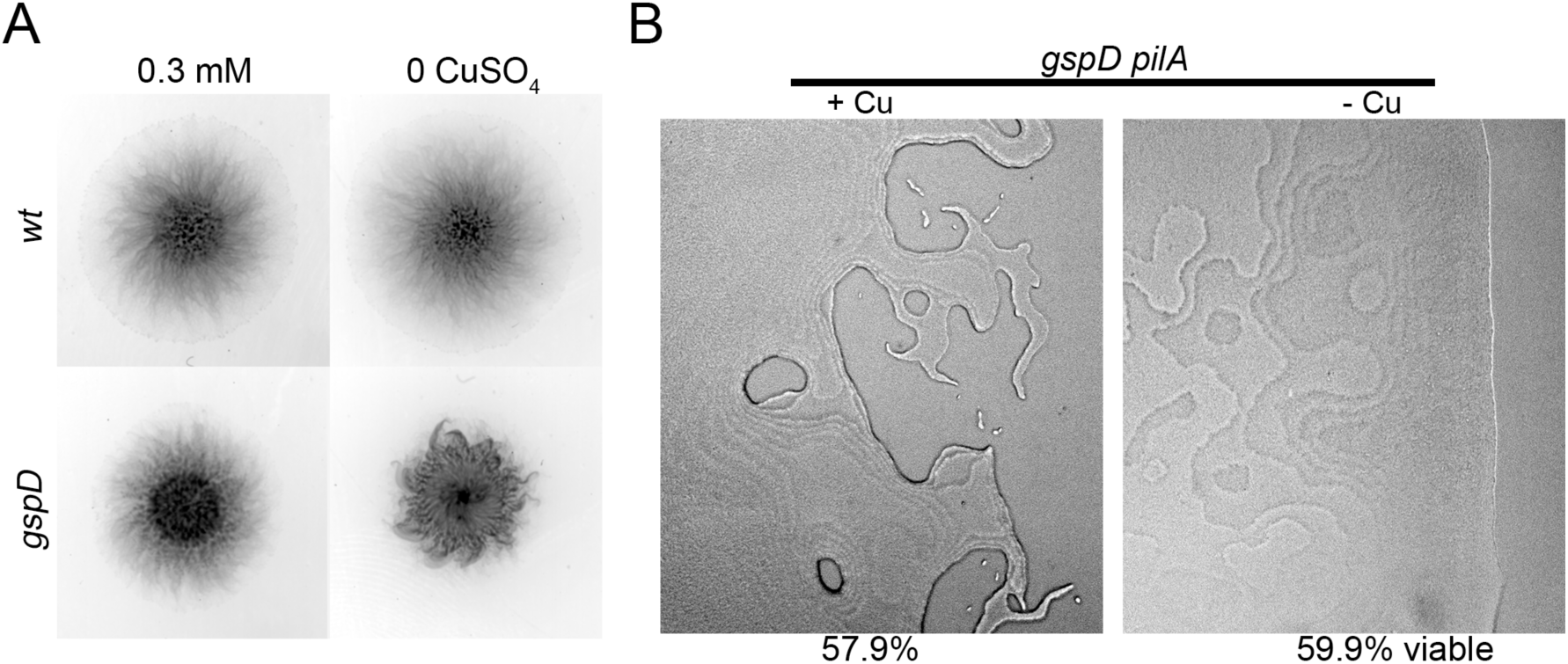
Swarming defect in GspD-depleted cells cannot be explained by failure of cells to grow. (A) To visualize social motility, *gspD* cells were grown in liquid culture in the absence of copper for 24 h, concentrated to 5 × 10^9^ cells/ml in CTT, and 10 μl were spotted onto 1/2× CTT plates with 0.4% agar, containing either 300 μM CuSO_4_ (0.3 mM) or 200 μM BCS (0 mM). Cells were allowed to swarm for 24 h and plates were scanned. All concentrations are mM CuSO_4_. (B) To visualize adventurous motility, *gspD ΔpilA* cells were grown in liquid culture in the absence of copper for 24 h, diluted to 1 × 10^8^ cells/ml in CTT, and 10 μl were spotted onto 1/2× CTT plates with 1.5% agar, containing either 100 μM CuSO_4_ (+ CuSO_4_) or 200 μM BCS (- CuSO_4_). After 48 h, cells were imaged by light microscope. To determine viability, cells were collected into PM buffer (20 mM Na-phosphate, 1 mM MgSO_4_, pH 7.4) and stained with the LIVE/DEAD *Bac*Light Bacterial Viability kit (Molecular Probes, Eugene, OR) according to the manufacturer’s instructions. Cells were imaged by fluorescence microscopy, and scored as viable (green) or dead (red) for each swarm (n > 400 cells); numbers indicate the percentage of viable cells.

**Figure S9.**
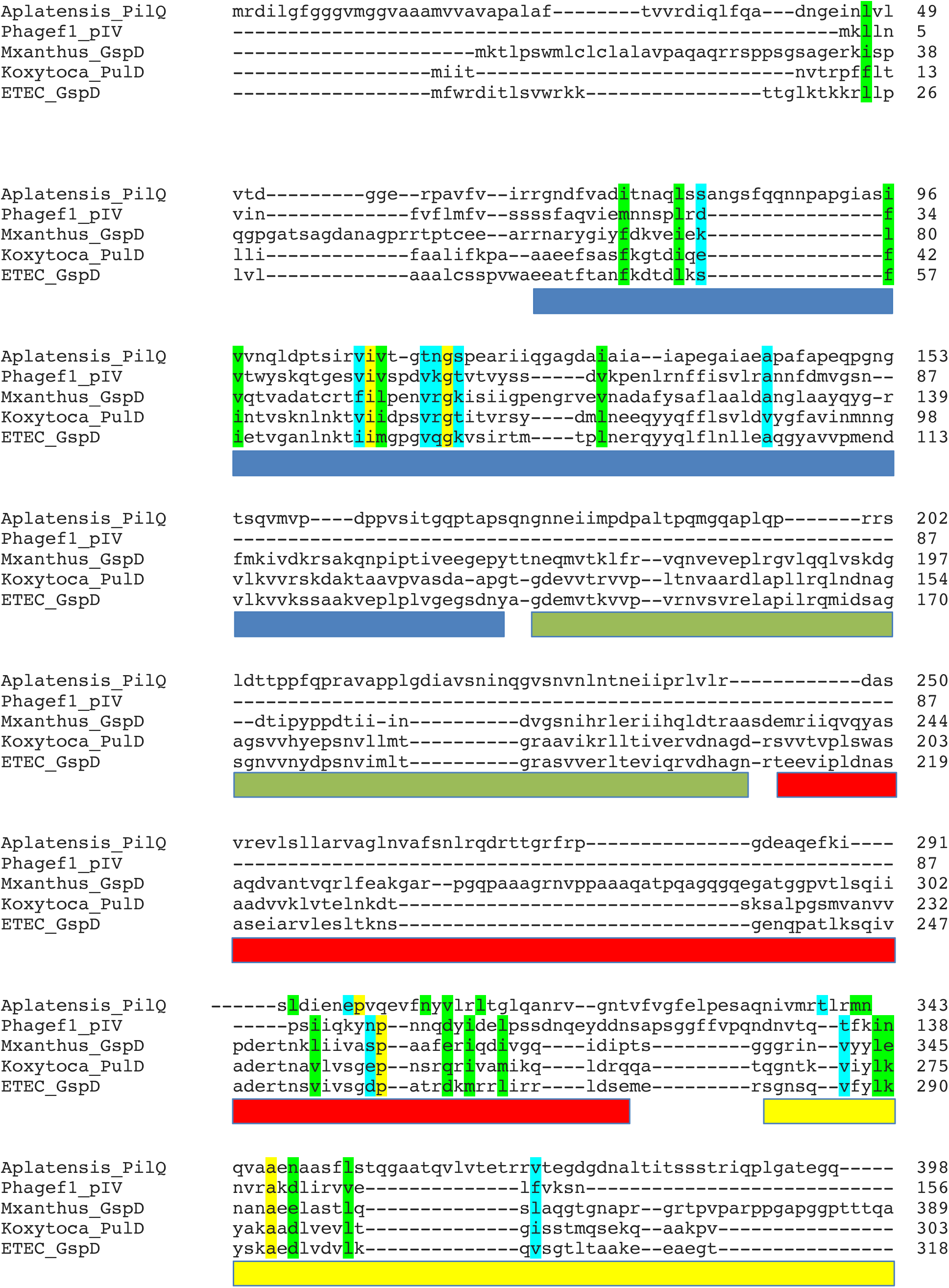
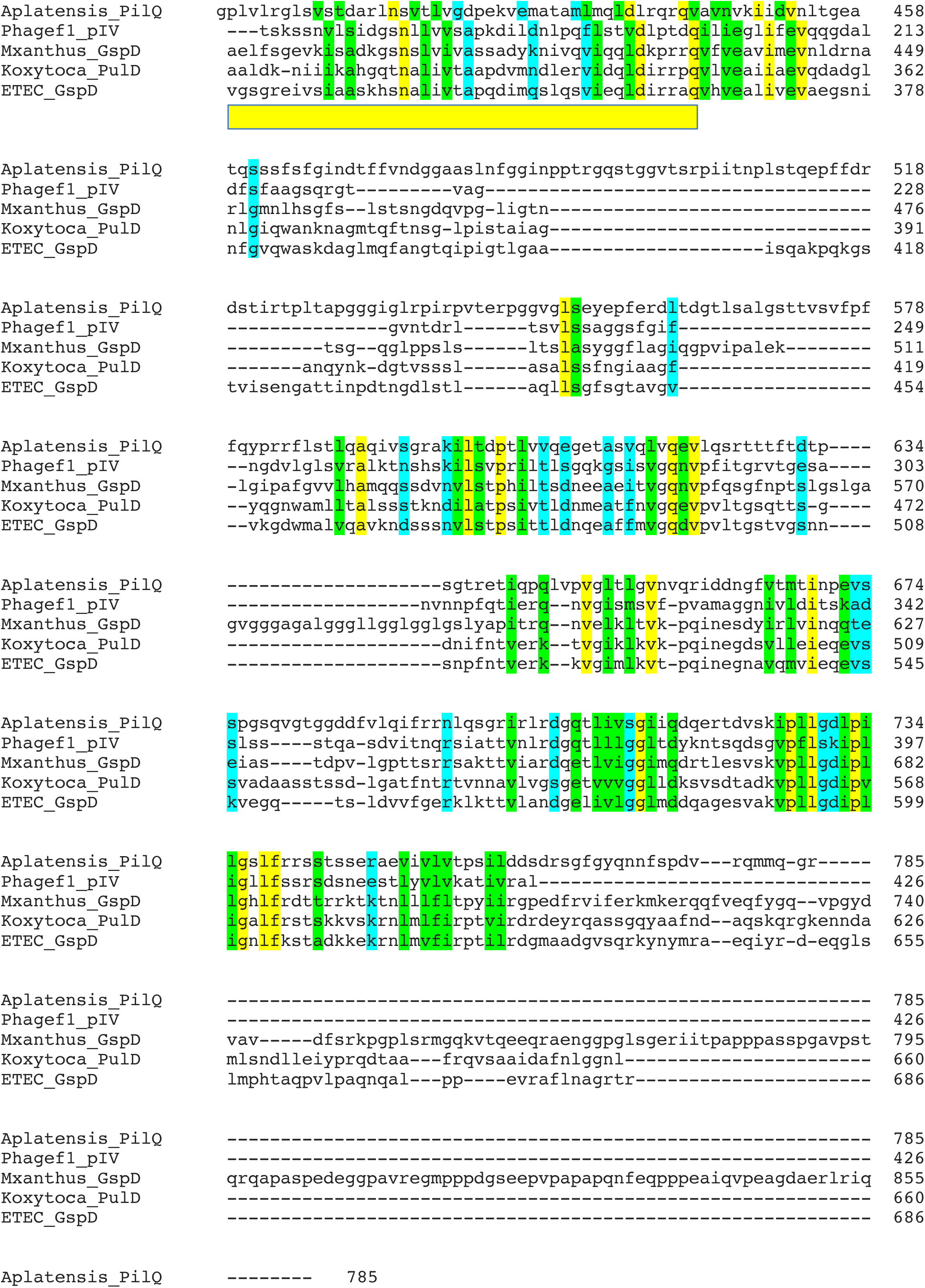
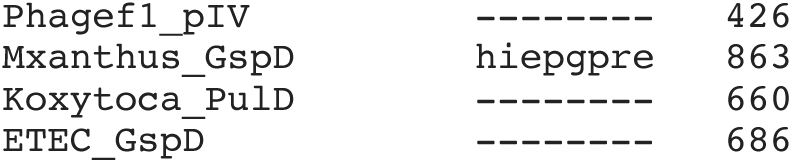
Multiple alignment of PilQ (*A. platensis*), pIV (phage F1), GspD (*M. xanthus*), PulD (*K. oxytoca*) and GspD (*E. coli* ETEC) using Clustal Omega. Yellow indicates positions occupied by single, fully conserved residues. Green positions indicate conservation between strongly similar residues that score > 0.5 in the Gonnet PAM 250 matrix, while blue indicates lower conservation scoring < 0.5 in the Gonnet PAM matrix. The color bars beneath the aligned sequences indicate the positions of the periplasmic N0 (light blue), N1 (green), N2 (red), and N3 (yellow) domains. All domains were assigned based on the published crystal structure of *E. coli* ETEC (23) and sequences were aligned using Clustal Omega (3).

**Table S1.** **Mass spec identification of the PilQ proteins of the various cyanobacterial species and of the GspD protein of *M. xanthus*.** Four independent isolations were used to identify PilQ of *A. platensis* using two different methods, Edman degradation and mass spectrometry, while GspD from *M. xanthus* was identified only by mass spectrometry. ^a^Note, that these mass spectrometry identifications were performed in 2011 at Harvard Mass Spectrometry and Proteomics Resource Laboratory, FAS Center for Systems Biology by microcapillary reverse-phase HPLC nano-electrospray tandem mass spectrometry (μLC/MS/MS) on a Thermo LTQ-Orbitrap mass spectrometer using MS/MS spectra and the MaxBP parameter for evaluation that are not any longer used. All other mass spectrometry measurements were done at the University of Sheffield using current methods described in the Materials and Methods section. *The M_w_ and the % sequence coverage represent the full-length protein. The number of aa is given with and without the signal peptide.

| Species | Type of Analyses | Mw* and aa | No. of peptides | Sequence coverage [%]* | Q-value | Score | MaxBP<sup>a</sup> | MS/MS Spectra<sup>a</sup> |
| --- | --- | --- | --- | --- | --- | --- | --- | --- |
| *Athrospira platensis* | Edman degradation | 81 kDa<br>785/757 | 2 totalling 26 aa | 3 | - | - | - | - |
|  | Mass spectrometry |  | 26 | 24 | - | - | 9.0e<sup>9</sup> | 647 |
|  |  |  | 21 | 20 | - | - | 7.2e<sup>9</sup> | 572 |
|  |  |  | 24 | 38 | 0 | 323 | - | - |
| *Phormidium autumnale* | Mass spectrometry | 82 kDa<br>805/778 | 29 | 46 | 0 | 323 | - | - |
| *Oscillatoria lutea* | Mass spectrometry | 83 kDa<br>810/783 | 158 | 72 | 0 | 323 | - | - |
| *Myxococcus xanthus* | Mass spectrometry | 90 kDa<br>863/842 | 29 | 40 | 0 | 323 | - | - |

## Notes

### Competing Interest Statement

The authors have declared no competing interest.

